# Cell Mechanics Regulate Membrane Tubulation-Driven Trogocytosis in Adherent Cells

**DOI:** 10.64898/2026.06.07.730655

**Authors:** Tanishka Agrawal, Binayak Banerjee, Shubham Padhi, Kishal Roy, Neha Paddillaya, Adhisha Roy, Santanu Talukder, Namrata Gundiah, Shivprasad Patil, Saroj K. Nandi, Sunando Datta

## Abstract

Trogocytosis is a widespread and physiologically important process in which one cell ingests fragments of another. It has been observed in diverse biological contexts, such as in *Entamoeba histolytica*, a human enteric parasite that nibbles host intestinal cells via trogocytosis, thereby invading the tissue and leading to lethal extra-intestinal disease, and in macrophages on interaction with adherent tumor monolayers. Despite its physiological relevance, knowledge of trogocytosis with adherent cells remains elusive. Here, we established a monolayer-based trogocytosis assay and uncovered a distinct membrane-tube-mediated trogocytic mechanism operative in adherent cells. We characterized it into four sequential events: contact, membrane tubulation, stretching, and scission. By targeting the proteic constituents of target cell stiffness, we show that trogocytic output exhibits a non-monotonic dependence on the viscoelastic properties of the target cell, with maximal uptake occurring in an intermediate mechanical regime. Further, we integrated live-cell trogocytosis observations with a viscoelastic model-based theoretical framework to propose a tube-breaking mechanism across distinct mechanical subtypes. Finally, using micropatterning, we demonstrated that target cell shape influences trogocytosis kinetics. This work suggests that trogocytosis is sensitive to the mechanical state of the target cell.

## Introduction

Trogocytosis (*from the Greek* trogo, *“to gnaw”)* was first introduced as the piecemeal ingestion of host cells by the pathogenic amoeba, *Naegleria fowleri*^1^. It is a contact-dependent process in which one cell physically nibbles and internalizes discrete fragments of another cell, in contrast to phagocytosis, which involves complete engulfment^2,3^. The functional consequence of Trogocytosis varies from a benign form of cell-cell interaction to cytotoxicity, depending on the context. In lymphocytes, it facilitates gentle intercellular communication through antigen acquisition without inducing cell death^4,5^, whereas immune effector cells such as Neutrophils perform intense trogocytosis to kill parasites^6^. Macrophages can similarly perform Fcγ receptor–mediated trogocytosis on antibody-opsonized tumor cells^7^. Beyond immunity, trogocytosis-like processes contribute to development and tissue remodeling, including synaptic pruning by microglia^8^ and cell remodeling during embryogenesis^9^. Trogocytosis also augments pathogenesis*. Entamoeba histolytica*, the causative agent of amoebiasis (>50 million cases annually),^10,11^ exploits trogocytosis to kill human cells^12^ and evade immune surveillance^13^. Previous studies suggest that upon intestinal colonization, the parasite may preferentially employ trogocytosis over phagocytosis to breach the epithelial barrier, progressively ingesting fragments of colonic epithelial cells and potentially facilitating dissemination to extraintestinal sites associated with severe disease^12^.

Given its broad biological relevance, it is important to study the factors that regulate this process. A preliminary study found that chemical stiffening of erythrocytes with glutaraldehyde reduced uptake by microphagocytosis (trogocytosis) in *E. histolytica*^14^, indicating that target cell deformability influences trogocytosis. Further studies in macrophages extended this principle by probing the role of target-cell rigidity in suspension systems. In particular, overexpression of Myocardin-related transcription factors (MRTFs) increased target cell stiffness (Young’s modulus) and reduced nibbling^15^. Recently, Fletcher et al. demonstrated that target cell cortical stiffness regulates the balance between phagocytosis and trogocytosis, with increased cortical tension suppressing trogocytic efficiency^16^. More recently, Ralston et al. showed that F-actin density in suspended Jurkat target cells modulates *E. histolytica* trogocytosis^17^.

Despite these advances, investigations into how target cell properties regulate trogocytosis have largely been confined to suspension systems. Although induced target cell adhesion in macrophage assays suggests that trogocytosis can predominate under adherent conditions^18^, how the physical properties of adherent cells govern this process remains unknown. These findings may contribute to improved strategies for combating parasitic infections.

In this study, we examined how the physical properties of adherent target cells influence amoebic trogocytosis. Additionally, employing adherent intestinal epithelial cells, the primary physiological targets of amoebic trophozoites, will improve the understanding of trogocytosis in the context of adherent systems. We established a novel monolayer-based trogocytosis assay that mimics physiological mechanical conditions to explore how the physical properties of adherent target cells affect trogocytic uptake. Using high spatial and temporal resolution, we identified a distinct mode of trogocytosis under this mechanical setting. Further using gene silencing, pharmacological modulation, and AFM, we probed the influence of the mechanical properties of the target cell on trogocytosis and observed a non-monotonic dependence of trogocytosis on the viscoelastic state of the cell. By integrating live-cell imaging data with a theoretical framework, we delineate the mechanistic differences in trogocytosis and the tube-breaking mechanism across different subtypes. Finally, using micropatterning, we showed that the target shape can modulate the kinetics of trogocytosis

## Results

### Trogocytosis involves Target Membrane Tubulation followed by Extension-driven Scission

Recent and earlier studies have demonstrated amoebic trogocytosis using suspension target cells^14,17^. Trogocytosis was observed on interaction with *ex vivo* mouse intestinal tissue^12^ suggesting *in vivo Entamoeba histolytica* trophozoites actively engage in trogocytosis of adherent intestinal epithelial cells^19^ (Fig. 1a). Therefore, to understand the mechanism of Trogocytosis under this mechanical setting, we developed a fluorescent monolayer-based trogocytosis assay that mimics the physiological mechanical environment, in which we used a membrane-labeled (WGA) adherent monolayer of cells as the target and amoebic trophozoites in suspension (Fig. 1b). We exploited two human gut epithelial cell lines - SW480 and DLD1 as the target cells. The trogocytic process was investigated at a sufficiently high spatial and temporal resolution using spinning-disk confocal microscopy. Unlike the suction-driven uptake mechanism for suspension cells^19^, we observed a distinct mode of Trogocytic events with adherent cells, which begins with the formation of a tubular protrusion on the host/target cell membrane post-adhesion, followed by its steady extension. Live-cell imaging also revealed that the amoeba moved away from the target cell during membrane extension, followed by scission of the pulled membrane tube predominantly near the amoeba interface, and the uptake of the broken fragments by the amoeba (Supplementary video 1-2). In some cases, the remaining membrane tube recoils back to the host cell surface (Supplementary video 3). Occasionally, we also observed retraction of the protruded membrane extensions without any scission, resulting in an unsuccessful event (Supplementary video 4). Based on our observations, we could describe a Trogocytic event as a process consisting of four key steps: (1) Contact; (2) Membrane protrusion Initiation; (3) Stretching; (4) Scission (Fig. 1c) (Supplementary Fig. 1a). Small, discrete WGA punctate observed inside the amoeba were treated as internalized cargo (Fig. 1b) and were utilized to quantify the efficiency of trogocytosis. In most of the cases (>90%), we also observed host actin in the extending membrane tube, when the host cells were labeled with the F-actin-specific fluorescent probe SiR Actin (Fig. 1d). Since the actin cytoskeleton is coupled with the host cell plasma membrane^20^, we observed a positive correlation (0.7) in intensity profiles of Actin and WGA across the stretched membrane tube (Fig. 1e). Normalized Actin intensity profiles along the membrane tubes onto which experimentally determined scission points were mapped revealed scission preferentially at regions of reduced actin intensity, while bright actin signals on the internalized piece were observed (Supplementary Fig. 1e) (Supplementary video 5). Previous studies measured trogocytosis efficiency as the percentage of recipient cells that acquired cargo^14–18^. Here, using a microscopy-based assay, we defined trogocytosis efficiency as the number of trogocytic events per amoeba, providing a direct measure of the magnitude of uptake. We counted WGA-positive fragments internalized by each amoeba. Since nuclei are not internalized by amoeba during trogocytosis, signals that colocalized with the nuclear marker Hoechst were excluded. The amoebic trophozoites displayed significantly higher trogocytic uptake for SW480 than DLD1, measured at multiple time points (Supplementary Fig. 1b-c). To better understand the kinetics of the process, we measured the rate of extension/stretching of protrusions, the maximum length of the protrusions/extensions prior to the scission, and the average width of the extensions. The dimensions of the membrane extensions were significantly different in the two cell lines we have studied (Fig. 1f). The SW480-originated membrane protrusions/extensions were significantly thicker and shorter compared to those measured in DLD1 (Fig. 1g) cells. We also observed a negative correlation (r= −0.4) between Maximum Length Stretched and the thickness of the membrane tube (Supplementary Fig. 1d). However, the extension rates measured for both cell lines were almost the same, with a value of approximately 0.24 μm/sec (Fig. 1h).

**Figure 1.**
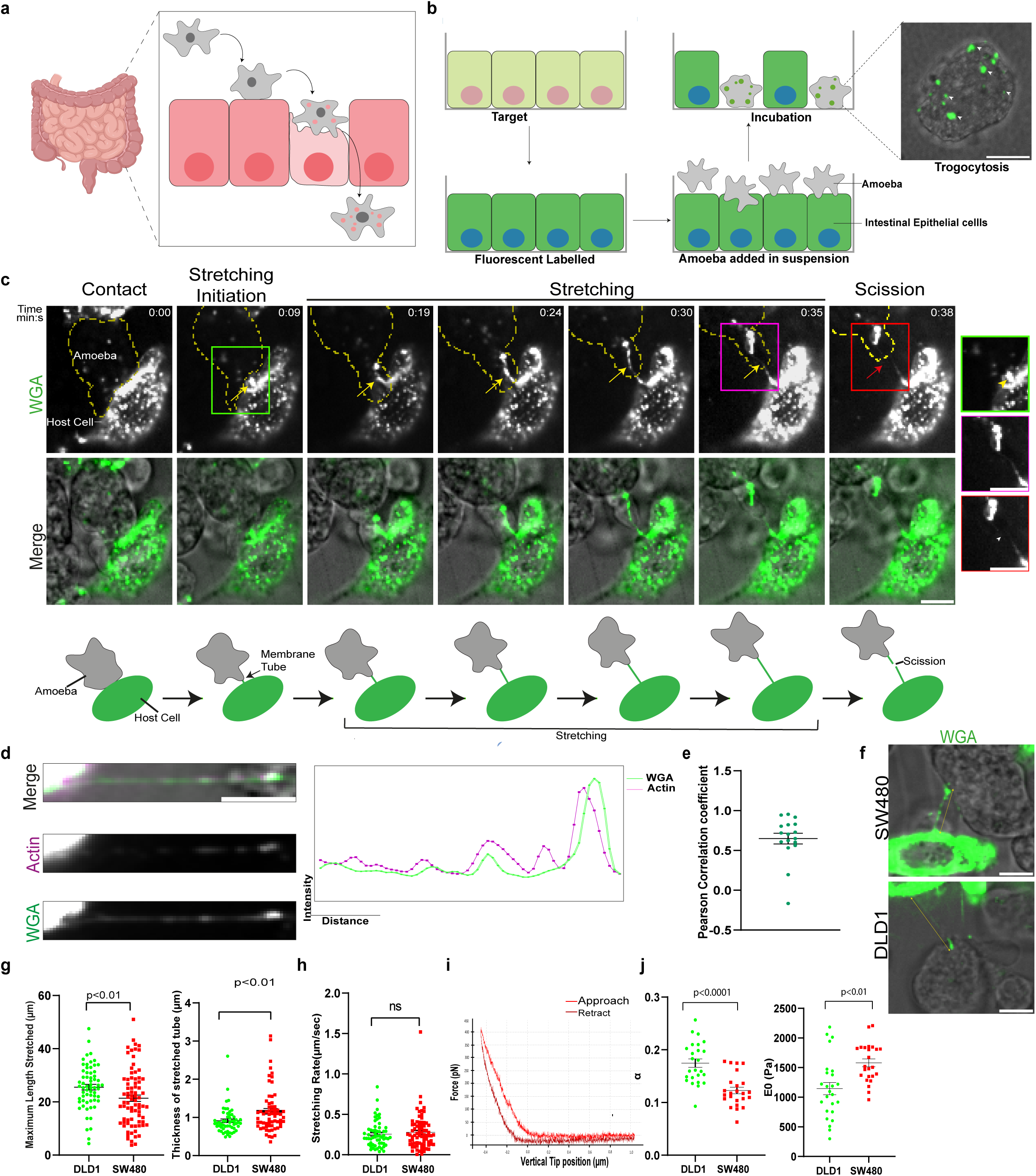
Trogocytosis Proceeds via Target Membrane Tubulation and Scission. (a) The *in vivo* model of amoebic trogocytosis. Motile amoeboid trophozoites colonize the large intestine, where they breach the epithelial barrier by trogocytosis of intestinal epithelial cells. (b) Schematic diagram of Monolayer-based trogocytosis assay. The adherent monolayer of colon cell lines was used as the target cells. The adherent monolayer of cells was membrane-stained using WGA (green) and Hoechst (blue) for the nucleus. The amoeba (grey) was added to the labeled monolayer in suspension, which then initiated trogocytosis. Trogocytosis is marked by small punctate bites of the host membrane fragments positive for WGA stain (Right). (c) Time-lapse montage showing a representative Trogocytic event. WGA-labeled adherent SW480 cells were presented to the amoeba, followed by time-lapse microscopy using a spinning disk confocal microscope. Frames were captured every 800ms. The fluorescence channel (WGA) and its merge with the bright field channel are shown. Scale bar 10µm. Trogocytic event is marked with a yellow arrow, and the red arrow indicates scission. The amoeba undergoing trogocytosis is outlined in yellow. Time is indicated as min: seconds in the top right of the fluorescence images, with time of contact defined as t = 0. Insets represent enlarged views of the regions enclosed by the respective coloured outlines. Inset yellow arrowhead marks the membrane tube, and the white arrowhead indicates the scission. Scale bar 10µm. (Bottom) Schematic representation of the trogocytosis process. (d) Host cells were labeled with a membrane stain (WGA) and an actin probe (SiR actin), and trogocytosis was imaged over time. Representative image of a pulled membrane tube by an amoeba in the Merge is shown. Magenta and green correspond to actin and the membrane, respectively. Scale bar 10µm. The membrane image as WGA channel and the actin channel are displayed individually below the merged image for clarity. Graph (right) represents the membrane and actin fluorescence intensity profile across the membrane tube. (e) Pearson correlation coefficient values for the intensity distributions of membrane and actin across the stretched tube just before scission were calculated. A total of 10 different extended membrane tubes were analyzed. (f) Representative image of extended membrane tube SW480 and DLD1 target cells labeled with WGA (green), where the yellow double-headed arrow corresponds to the stretched membrane tube. Scale bar 10µm. (g) Quantification of the length of the extended membrane tube just before scission (left) and the thickness of the membrane tube (Right) for SW480 and DLD1 cells. Each data point represents a dimension value derived from a single Trogocytic event. Mean and error bars (denoting SEM) are indicated in black. A total of 62 trogocytic events from 10 independent experiments were analyzed for DLD1 cells, and 79 trogocytic events from 12 independent experiments were analyzed for SW480 cells. (h) Quantification of the membrane stretching rate for DLD1 and SW480 target cells by the amoeba. Each data point represents a value obtained by tracking a single Trogocytic event, where n=59 for both cells. Mean and error bars (denoting SEM) are indicated in black. (i) A typical force curve obtained from a live SW480 cell during atomic force microscopy probing is shown, with the green line representing the approach curve and the red line representing the retraction curve. (j) Quantification of the power-law exponent (α) and the instantaneous Young’s modulus (E_0_) derived from AFM measurements for SW480 and DLD1 cells with n=26 and 25 cells analyzed respectively. In the graph, each data point represents the average of values obtained from 12 best-fit curves per cell. Mean values are shown with black lines, and error bars represent the SEM. Statistical significance was assessed for all the data using an unpaired t-test.

We hypothesized that in DLD1 and SW480 cells, membrane protrusions are stretched to varying degrees prior to scission, possibly due to differences in target cell properties, and one of the potential factors could be the target cell deformability. Cell mechanics are often quantified using Young’s modulus; however, cells display both elastic and viscous behavior^21^. Accordingly, we characterized their viscoelastic properties using atomic force microscopy (AFM). Cells were probed at the center using an AFM cantilever attached with a spherical bead with a diameter of 5-µm. For each condition, AFM force-indentation curves were obtained (Fig. 1i). We then determined the instantaneous Young’s modulus and the power-law exponent (α) by analyzing the force–indentation curves using Ting’s model^22,23^. E_0_ represents the sample’s rigidity, whereas the power-law exponent α is a measure of the sample’s fluidity and dissipative properties^21^, which were used to describe the viscoelastic state of the cell. We found that DLD1 cells displayed 1.4-fold higher α and 1.37-fold lower E_0_ values than SW480, thereby exhibiting relatively soft, fluid-like behavior (Fig. 1j). These findings suggest that the properties of target cells may influence the overall efficiency of the trogocytic process in the parasite.

### Amoebic Trogocytosis is Responsive to perturbations in the Target Cell actin cytoskeleton

To establish that the cell mechanical properties indeed influence the amoebic trogocytic efficiency, we altered the viscoelastic properties of the target cell by perturbing its actin architecture^24–27^ and conducted time-lapse imaging of the trogocytic process under physiological conditions, as well as measured the average trogocytic efficiency of the amoebic trophozoites. We treated the target cell monolayer with four different actin modulators: Latrunculin A and Cytochalasin D, which promote actin depolymerization; Jasplakinolide, an actin stabilizer; and CK869, an inhibitor of the Arp2/3 complex^28^ involved in actin nucleation^29^. We first confirmed their effect by studying F-actin structures in the drug-treated cells. We observed that both Latrunculin A^30^ (Fig. 2a) and Cytochalasin D disrupted the actin network and led to the formation of actin foci^31^ (Supplementary Fig. 2b), whereas Jasplakinolide reorganized the actin cytoskeleton into amorphous aggregates^32^(Fig. 2b). In the case of CK869 we observed the formation of blebs^33^ (Fig. 2c). To evaluate the effect of target cell stiffness on trogocytosis, we presented drug-treated target cell monolayers to the amoebic trophozoites and allowed trogocytosis to occur; we then quantified differences in their trogocytic uptake. Target cells treated with an equivalent amount of DMSO to that of the drug were used as a control. We observed a 4.2-fold reduction in trogocytic uptake in Latrunculin A-treated cells (Fig. 2d) and an enhancement of trogocytosis (1.2-fold) in Jasplakinolide-treated target cells compared to control (Fig. 2e). We also observed a 1.4-fold increase in Trogocytic uptake upon treatment with CK869 (Fig. 2f). However, Cytochalasin D treatment did not result in any significant difference in trogocytosis efficiency (Supplementary Fig. 2a) due to its high K_off_ of binding to actin in, allowing cells to recover quickly upon its removal (Supplementary Fig. 2b). In contrast, the effect of Latrunculin A, Jasplakinolide, and CK869 persists after removal (Supplementary Fig. 2b). Next, to address whether the observed extent of trogocytosis with the drug-treated target cells is associated with the latter’s viscoelastic properties, we measured their mechanical properties by probing them with Atomic Force Microscopy post-pharmacological interventions. We found that Latrunculin A treatment markedly altered the viscoelastic state of the cells, resulting in a softer and more fluid-like mechanical phenotype characterized by an increase in the α from 0.17 in control cells to 0.23, accompanied by a reduction in the instantaneous Young’s modulus (E_0_) from 1940 Pa to 794 Pa whereas, Jasplakinolide treatment led to α value 0.2399, while maintaining an E_0_ to that of control cells 1707 Pa (Fig. 2g-h). We observed a similar trend in DLD1 cells (Supplementary Fig. 2c). Together these observation suggest that amoebic trogocytosis is responsive to the material properties of target cells and this relationship does not appear to follow a simple monotonic trend.

**Figure 2.**
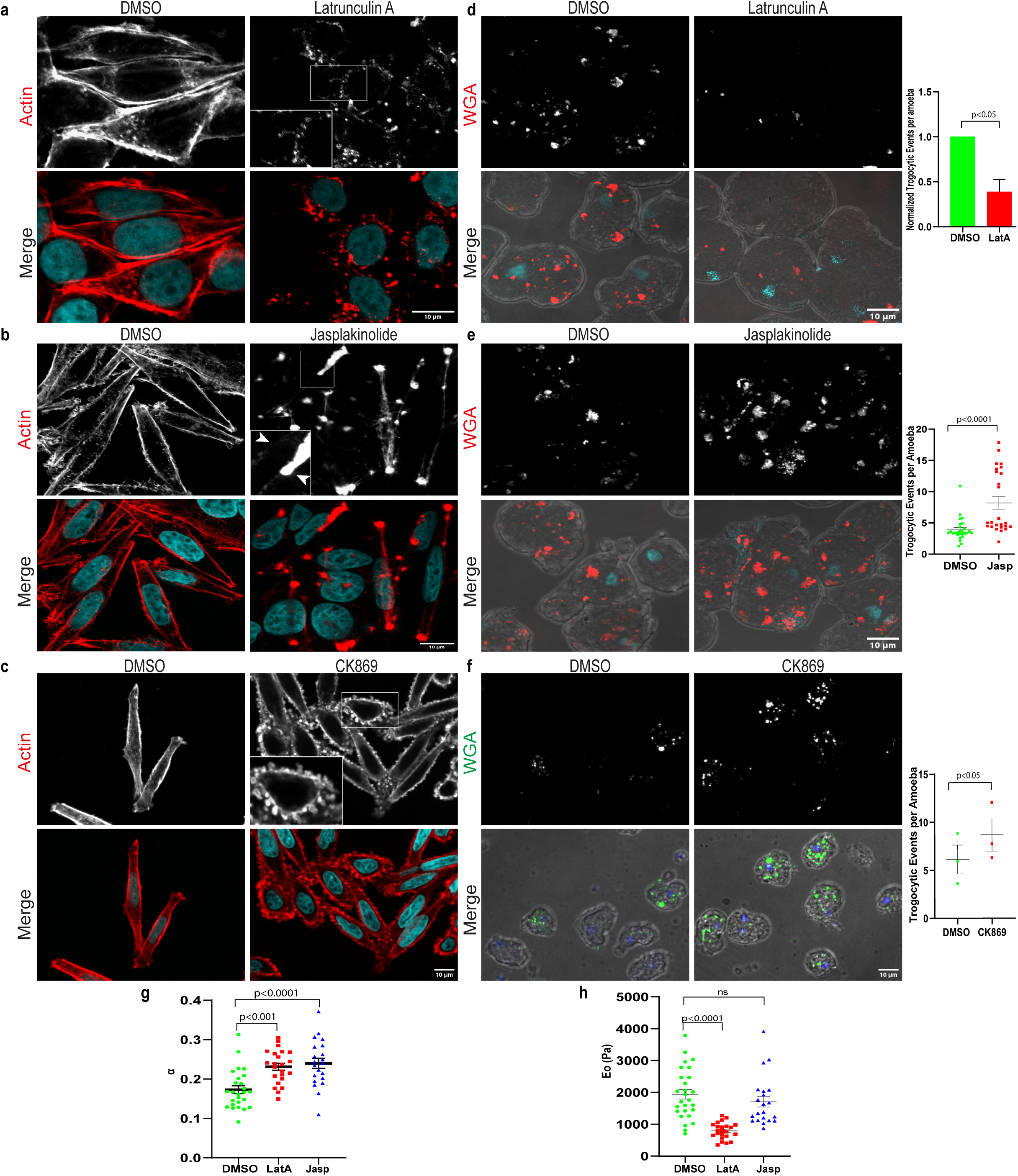
Target Cell Actin Perturbations Modulate Amoebic Trogocytosis. (a) The images show the actin staining for host cells that were treated with actin depolymerizer Latrunculin A (1µM) and an equivalent amount of DMSO for 2 h. After treatment, the cells were fixed, permeabilized, and stained with Phalloidin 568 to visualize actin architecture and DAPI. The images show actin staining (Red) and DAPI (cyan) in DMSO-treated and Latrunculin-treated SW480 cells, visualized by confocal microscopy. Scale bar 10µm. Insets show the enlarged view of the region inside the box. (b) Host cells were treated with 0.2 µM Jasplakinolide, an actin stabilizer, for 1 h. Cells were treated with an equivalent amount of DMSO as a control. Cells were fixed, permeabilized, and stained with Phalloidin 568 and DAPI. The images were captured using confocal microscopy. The actin channel (red) and the merged image, which shows both actin and DAPI (cyan), are shown. Scale bar 10µm. Insets show the magnified view of the region inside the box. The white arrow marks the differences in the cortical actin distribution. (c) Actin staining for Host cells treated with DMSO and 40µM CK869 for 1h. Phalloidin 568 is shown in red, and the merge shows actin and DAPI (cyan) channels. Scale bar 10µm. Insets show the enlarged view of the region inside the box. (d) Representative images for Trogocytosis. Images shown are a Maximum Intensity Projection (MIP). The SW480 cells, after treatment with Latrunculin A for 2h and DMSO, were first stained with membrane stain WGA and the nuclear stain Hoechst, incubated with amoeba for 40 minutes, and then fixed. The images were then captured in confocal microscopy. The image shows the WGA channel (Red) and its merge with DIC to visualize the amoeba. Scale bar 10µm. Small WGA puncta are visible within the amoeba due to trogocytosis. (Right) Microscopy-based analysis of Trogocytosis efficiency, defined as the total number of Trogocytic events normalized by the total number of amoeba in one field of view. For each independent experiment, the average value was calculated and normalized to the corresponding DMSO control. Each data point represents the normalized mean from an independent experiment. The data were obtained from three independent experiments, and a total of 679 amoeba were analyzed. p-values were determined by using a paired t-test. (e) Representative trogocytosis images for DMSO and Jasplakinolide-treated cells. Images shown are a Maximum Intensity Projection (MIP). Top: WGA channel (Red) to show internalized fragments inside the amoeba, Bottom: WGA channel merged with DIC to visualize the amoeba. Scale bar 10µm. (Right) Quantification of trogocytosis efficiency was analyzed as described above. N=3 independent experiments are pooled together. Each data point corresponds to a single frame, n=1202 amoeba for both conditions. Mean and SEM are indicated with black lines. (f) Representative images of trogocytosis with SW480 cells stained with WGA after treatment with DMSO and CK869 for 1h, and coincubated with amoeba for 15 minutes. Shown are the WGA channel (Green) and the Merge channel with DIC. Images shown are a Maximum Intensity Projection (MIP). Scale bar 10µm. (Right) Microscopy-based analysis of trogocytosis efficiency. Each point represents the mean value from an independent experiment with n=451 amoeba and N=3. Horizontal black lines represent the overall mean and error bars (denoting SEM). The paired t-test was used to determine the statistical significance. (g-h) Quantification of Power law component (α) (g) and Instantaneous Young’s modulus (E_0_) (h) determined from AFM. For each condition, at least 22 cells were analyzed. The mean and error bars (denoting SEM) are represented with black lines. The p-values for both plots were determined using a one-way ANOVA with Dunnett’s multiple comparisons test.

### Relationship between Trogocytosis and Actomyosin-based Cell Stiffness

The organization of actin filaments and their interplay with actin crosslinkers and Myosin motors determine the stiffness of the cell^34^. We first inhibited Myosin II in SW480 cells using Blebbistatin, which prevents myosin binding to actin. We observed no significant change in Trogocytosis uptake (Fig. 3a-b) and no alteration in α and E_0_ parameters as derived by AFM (Fig. 3c). We also observed minimal perturbation of the actin architecture, as evident by actin staining, upon myosin inhibition (Supplementary Fig. 3a). This suggests that, in this cell line, stiffness is not tuned by Myosin activity^35^; hence, trogocytosis efficiency remains unaffected. Therefore, we assessed trogocytosis efficiency in DLD1, which exhibits a more organized stress fiber network. Stress fibers arise from actomyosin contractility^36,37^ studies have established a positive correlation between stress fiber abundance and cellular stiffness^38,39^. Accordingly, we hypothesized that modulation in stress fiber levels would influence the stiffness of the cell. Upon Myosin II inhibition with Blebbistatin in this cell line, we observed a significant loss of stress fibers on Blebbistatin treatment (Supplementary Fig. 3b), which corroborated with increased trogocytosis efficiency (2-fold) (Fig. 3d-e). Given the established contribution of stress fibers to global cell stiffness, these findings suggest that enhanced trogocytosis may arise from reduced cellular elasticity following stress fiber disruption. We further confirmed our observation by depleting Myosin II in DLD1 cells (Supplementary Fig. 3c). A similarly disrupted stress fiber network was also observed upon Myosin II knockdown (Supplementary Fig. 3d). We then performed Trogocytic assay with host cells with depleted levels of Myosin II, and we observed a 2-fold increase in amoebic trogocytosis uptake (Fig. 3g), a similar extent observed with blebbistatin. We next measured the mechanical properties of cells following Blebbistatin treatment using AFM. Blebbistatin treatment resulted in a modest increase in the fluidity index (α) accompanied by a slight reduction in the instantaneous Young’s modulus (E_0_) (Fig. 3f). This discrepancy with previous studies linking stress fibers to stiffness could be attributed to differences in AFM probe geometry and model used^38,39^.

**Figure 3.**
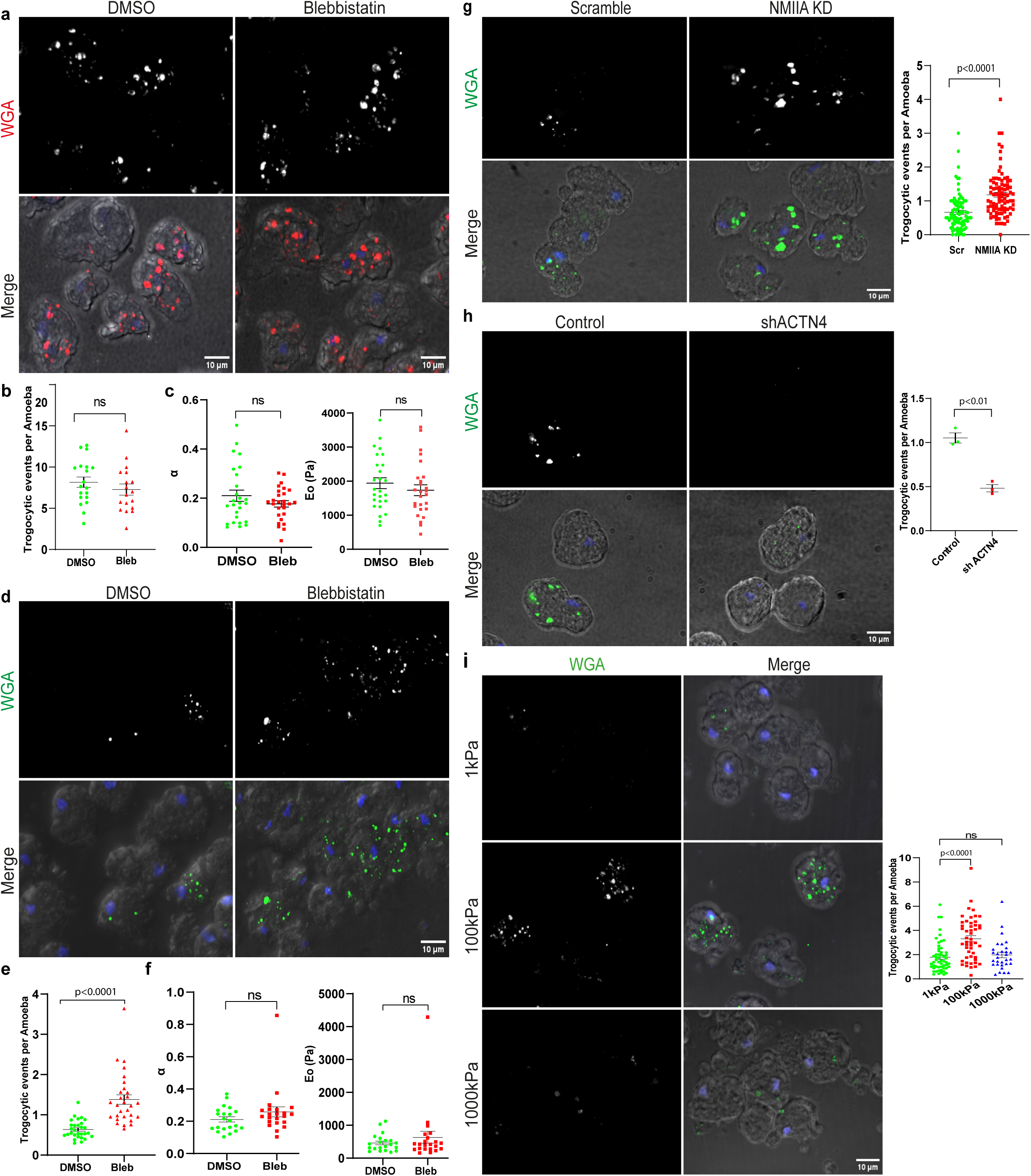
Relationship between Trogocytosis and Actomyosin-based Cell Stiffness. (a) Representative images of trogocytosis with SW480 cells treated with 75 µM Blebbistatin or DMSO for 4h, and were labeled with WGA and Hoechst, incubated with amoeba for 40 min, fixed, and visualized by confocal microscopy. The images show the WGA channel (Red) and the Merge channel with DIC. Images shown are a Maximum Intensity Projection (MIP). Scale bar 10µm. (b) Quantification of trogocytosis efficiency, expressed as the number of trogocytic events normalized to the total number of amoeba per field of view. Each point represents a value from a single field of view. N=2 independent experiments pooled together, with n=880amoeba. The black lines represent the mean and SEM. Statistical significance was evaluated using an unpaired t-test. (c) Quantification of the power-law exponent (α) and instantaneous Young’s modulus (E_0_) obtained from AFM analysis. Each data point represents the mean of 12 best-fit curves for an individual cell, with n = 26 cells analyzed per condition. The mean values and SEM are indicated by black lines. The p-values for both plots were determined using an unpaired t-test. (d) Representative images of trogocytosis with the WGA (Green) channel and the Merge channel with DIC are shown. DLD1 cells were treated with 75 µM Blebbistatin or DMSO for 4 h, labeled with WGA and Hoechst, incubated with amoeba for 25 minutes, fixed, and imaged using confocal microscopy. Images shown are the Maximum Intensity Projection (MIP). Scale bar 10µm. (e) Quantification of Trogocytosis efficiency as the number of trogocytic events normalized to the total number of amoeba per field of view; each point represents a trogocytosis efficiency value from a single field of view. Data were pooled from three independent experiments; n = 2000 amoeba. (f) Quantification of α and E₀ parameters derived from AFM. Each data point represents the average value from the 12 best-fit curves for each cell, with n ∼ 21 per condition. (e-f) The mean and SEM are represented with black lines. All p-values were determined using Student’s unpaired t-test. (g) Representative trogocytosis images showing WGA signal inside the amoeba after trogocytosis with WGA and Hoechst labeled control (transfected with scrambled siRNA) and NMIIA knockdown DLD1 cells for 30minutes. The MIP with the WGA channel (Green) and the merge channel with DIC are displayed. Scale bar 10µm. (Right) Trogocytosis efficiency for Scramble and NMIIA knockdown DLD1 cells. Each data point value is from a single field of view. Data were pooled from four independent experiments; n = 3000 amoeba. The black bars represent the mean and error bars (denoting SEM). The statistical significance was assessed using an unpaired t-test. (h) Representative images of trogocytosis with control and α-actinin 4 knockdown DLD1 cells are shown. DLD1 cells following 70h of transfection were labeled with WGA and Hoechst and challenged with amoeba for 30 minutes. Shown are the WGA channel (Green) and the merge channel with DIC. Images are Maximum Intensity Projections (MIPs). Scale bar 10µm. (Below) Trogocytosis efficiency is quantified as the number of trogocytic events internalized per amoeba. Each data point represents the mean value from a single experiment. Horizontal black lines represent the overall mean and error bars (denoting SEM); n = 2000 amoeba and N=3. The p-value was determined using a paired t-test. (i) Representative Trogocytosis MIP images for DLD1 cells cultured on three different polyacrylamide substrates (1kPa), (100kPa), and (1000kPa). After 24h of cell seeding, the cells were stained with WGA and Hoechst, incubated with the amoeba for 35 minutes, and analyzed by confocal microscopy. The images show the WGA channel (Green) to indicate the Trogocytic signal inside the amoeba and the merge channel with DIC. (Bottom) Trogocytic efficiency graphed against substrate stiffness. The black lines indicate the mean and SEM. N=3, n=1197 amoeba for 1kPa and 100kPa and N=2, n= 602 amoeba for 1000kPa. The p-value was obtained using a one-way ANOVA test with Dunnett’s multiple comparisons test.

Actin cross-linkers, the other class of molecules that influence cell stiffness, we set out to alter the expression of one of these cross-linkers and assessed its effect on amoebic trogocytic efficiency. We chose Alpha-actinin 4 because it has previously been shown to have an opposing effect on stress fiber stability^40,41^. Consistent with this, depleting the level of ACTN-4 led to an increase in the number of stress fibers (Supplementary Fig. 3e-f), suggesting increased cell stiffness. Next, to determine how it impacts amoebic trogocytosis, we incubated amoebae with ACTN4 knockdown cells and control cells and observed approximately a 2-fold decrease in trogocytosis (Fig. 3h). Together, these findings suggest that increased cellular stiffness associated with elevated stress fiber levels may negatively regulate trogocytosis.

To further confirm that host cell stiffness influences trogocytosis, we employed a polyacrylamide gel of varied stiffness as a substrate to tune cell mechanics. Previous studies have highlighted how physical input from the underlying substrate can reinforce cytoskeletal reorganization into more ordered actin bundles and, ultimately, alter the mechanical properties of the cell, with cell stiffness increasing as a function of substrate stiffness^42,43^. We analyzed amoebic Trogocytic efficiency for host cells cultured on Polyacrylamide gels of three different Young’s moduli: 1kPa (soft), 100kPa (intermediate) and 1000kPa (Hard)^44^ (Supplementary Fig. 3g). We observed a ∼1.9-fold higher trogocytic uptake in cells cultured on an intermediate substrate than with cells on a soft substrate, and a ∼1.7-fold higher uptake than with cells on a hard substrate. (Fig. 3i). This experiment further confirms that Trogocytic efficiency is influenced by the factors that alter cell mechanical properties via modulation of the actin cytoskeleton assembly.

### Theoretical Model for Trogocytic Membrane Remodeling

Our experimental data show four distinct stages of the trogocytosis process: (1) contact formation, (2) protrusion initiation, (3) stretching, and (4) scission. To understand the dynamics of the process and the underlying detailed mechanism, we have developed a mechanistic model of the last two stages, namely the stretching of the protrusion and its scission. We model the protrusion comprising n_0_ elementary units. These units have their own mechanical properties, which can vary for different units; however, for simplicity, we assume they are the same within a protrusion. Furthermore, these properties will depend on the mechanical state of the cell, governed by the visco-elastic properties of the cell cortex, and can be affected by cell shape. Within the model, these cellular properties enter via the mechanical properties of the individual units. The amoeba attaches to these units via specific ligands and pulls the entire protrusion until it ruptures, leading to the scission event. To account for the viscoelastic nature of the system, we model each unit via a Kelvin-Voigt (KV) model. A KV unit consists of a spring with spring constant k and a dashpot with viscosity ξ connected in parallel. The physical state of the cell will affect both k and η. As shown schematically in Fig. 4a, the entire protrusion consists of n_0_ such KV units. We can then calculate (see Supplementary Notes) the instantaneous protrusion length as ℓ(t) = ℓ_0_ + a(1 − e^−bt^), where we have measured time *t* from the beginning of our measurement process, when ℓ(t=0) = ℓ_0_. The constants are related to the protrusion thickness and mechanical properties of the system: a = F_0_/n_0_k and b = k/ξ, where F_0_ is the total force with which the amoeba pulls the particular protrusion, assumed constant during the process, and *ξ* is the viscous force.. In our experiments, we have measured the instantaneous lengths, ℓ(t), as a function of t (Fig. 4b). Sometimes, there is a lag before the amoeba starts pulling the protrusion; this lag time is possibly related to the biochemistry of the process. We have excluded this lag time from our analysis. We first show that the experimental data are consistent with the hypothesis of the theory. For this, we fit our model with ℓ_0_, a, and b as fitting parameters. If we plot (ℓ(t) − ℓ_0_)/a as a function of bt, we expect data collapse to a master curve as shown in Fig. 4c. From the fits, we obtain an average value ⟨b⟩ = 0.035*s^-1^*; this is the typical pulling rate in the experiments. The excellent data collapse in Fig. 4c supports the validity of the model.

**Figure 4.**
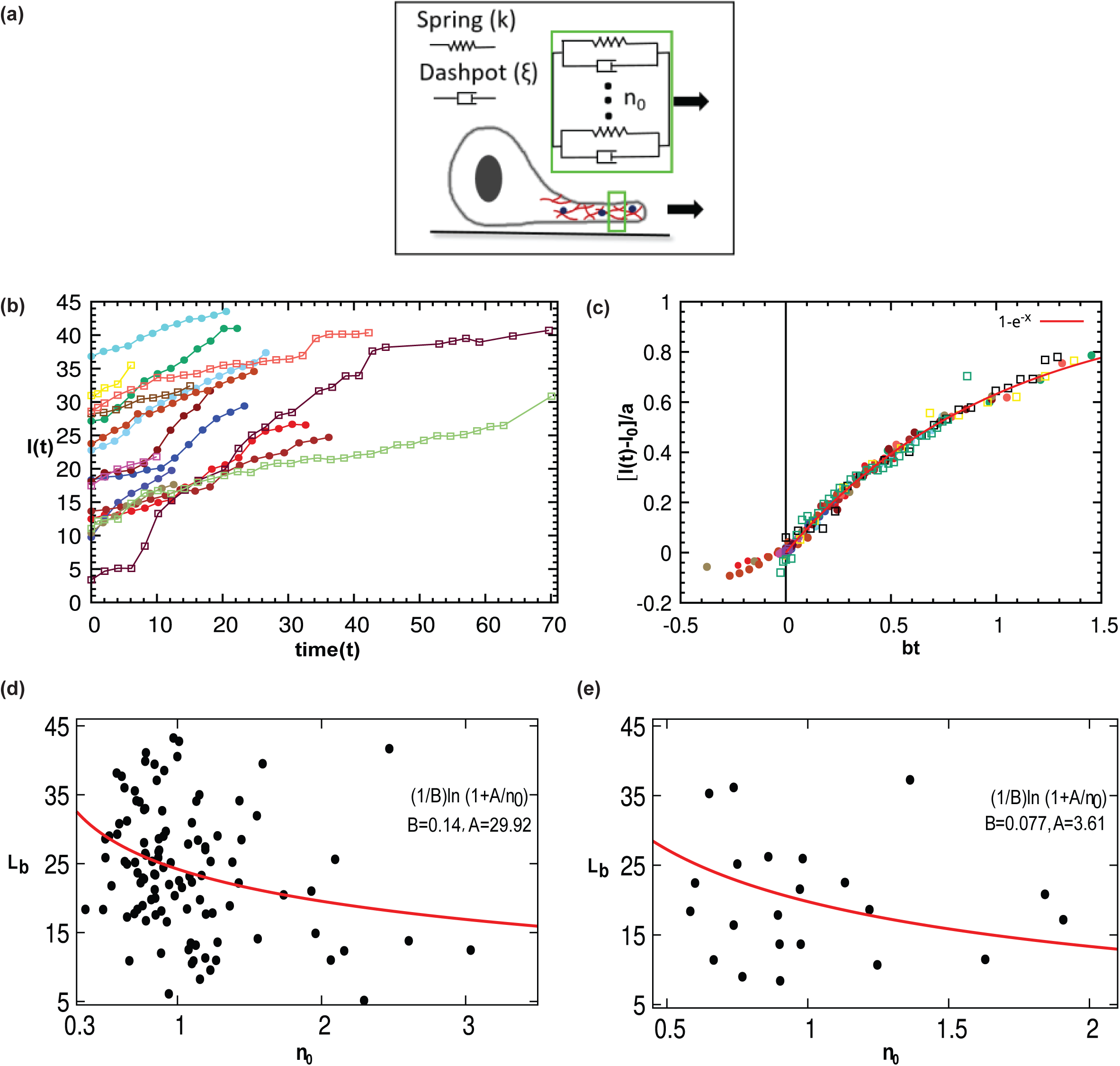
Theoretical model for Trogocytosis dynamics. (a) Schematic of the model in which the protrusion is composed of parallel Kelvin-Voigt units. (b) Experimental wild-type data showing the variation of extension length with time (dots: SW480; boxes: DLD1). The extension lengths, l(t) are fitted to obtain the a, b, and l_0_. (c) Upon rescaling time bt and length (l(t) − l_0_)/a, all curves collapse onto a single master curve, (1 − *e*^−*x*^). Negative bt indicates the lag period before pulling. Experimental data of breaking length as a function of initial thickness for (d) WT and (e) Latrunculin-treated cell lines (SW480 and DLD1 datasets combined). The solid line represents the theoretical predictions (see SM).

Given this microscopic picture, we now focus on the breaking mechanism. As shown in Supplementary Fig. 1d, the breaking length, L_b_, decreases monotonically with protrusion thickness. This is different from the typical behavior of highly viscous fluid, where surface tension dominates, and a thicker object typically extends further before breaking. The data suggests failure of individual elements dominates the breaking of the protrusion via a nucleation like mechanism. Considering the extension rate constant (∼ 0.24μm/sec, Fig. 1h), we can write the failure rate of a single element as r(ℓ) = r_0_e^αℓ^, with r_0_ and α being constants. Then the survival probability of all the elements becomes 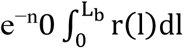. When an amoeba pulls the protrusion, if one of the elements breaks, the surviving elements must balance the same force, increasing the average force on each of the elements. This exponentially increases the probability of breaking for the remaining elements. Considering this as a fast process, we can obtain the breaking length as a function of n_0_ as L_b_ = ln[1 + A/n_0_]/B, where A and B are constants. As shown in Fig. 4d, the theory captures the general trend of the experimental data for L_b_. Note that both A and B depend on the mechanical properties (Supplementary Notes) of the individual cells leading to the variability of L_b_. In addition, we can also calculate the protrusion breaking time, T_b_, from the theory. As shown in Supplementary Notes Table I, the T_b_, is smaller for the WT SW480 cells compared to the DLD1 cells. This comes from the smaller stretching rate for the latter.

Furthermore, the theory shows that the scission mechanism remains the same for the Latrunculin-treated cells as well. We show in the Supplementary Notes that the Latrunculin A cells also follow the theory, with the individual experimental data collapsing onto a master curve. The qualitative behavior of L_b_ as a function of protrusion thickness remains the same as in the WT case and agrees well with the theory (Fig. 4e). In addition, as Table II in the Supplementary Notes shows, the protrusion breaking time, T_b_, nearly doubles for the Latrunculin A-treated cells. This explains why these cells, despite being mechanically softer, shows fewer trogocytic events in the experiment.

### Trogocytosis dynamics depend on the target cell Shape

Within the human host*, Entamoeba histolytica* encounters target cells with varying physical properties, such as stiffness and shape. However, it is yet to be determined whether the target cell shape influences amoebic trogocytosis. Here, we set out to decipher the influence of target cell shape on trogocytosis. We used ECM-coated dishes to direct cell shape change. ECM binding to integrin can initiate cytoskeletal organization, increasing its aspect ratio^45^. We performed the Trogocytic assay with cells cultured on collagen-coated (25 ug/ml) and uncoated dishes. Cells adopted an elliptical morphology (Aspect ratio 4-5) on a collagen-coated surface, while they remained largely round-shaped on glass (Supplementary Fig. 4a). On quantification of Trogocytic efficiency, we observed a 1.35-fold increase in trogocytosis for elliptical host cells on collagen compared to round cells on glass (Fig. 5a and 5c). We also performed a similar assay with DLD1 cells (Supplementary Fig. 4b) and observed a 1.62-fold increase in trogocytic efficiency (Fig. 5b and 5c). The above results suggest that target cell shape may be an important factor in determining trogocytic efficiency. However, to rule out any possible contributions from collagen, we employed micropatterning (Supplementary Fig. 4c), a novel tool for tuning cell shape. Cells were confined to grow on either circular (diameter ∼22 µm) or elliptical (aspect ratio∼4) collagen micropatterns (Fig. 5d) with almost the same surface area (Fig. 5e) imprinted with PDMS stamps. We performed time-lapse imaging under physiological conditions using spinning disc confocal microscopy to study the kinetics of amoebic trogocytosis on these patterns. We then extracted kinetics information from a single Trogocytic event for both target shapes (Fig. 5f) (Supplementary Videos 6-7). We observed that the interval between contact and stretching initiation is significantly longer for circular cells compared to elliptical cells, demonstrating a clear influence of target cell shape on the amoebic trogocytosis. In addition to this, in more than 90% cases, we also observed that membrane tube initiation preferentially occurred from the major axis in elliptical cells. These results highlight the contribution of target cell shape in regulating trogocytosis kinetics.

**Figure 5.**
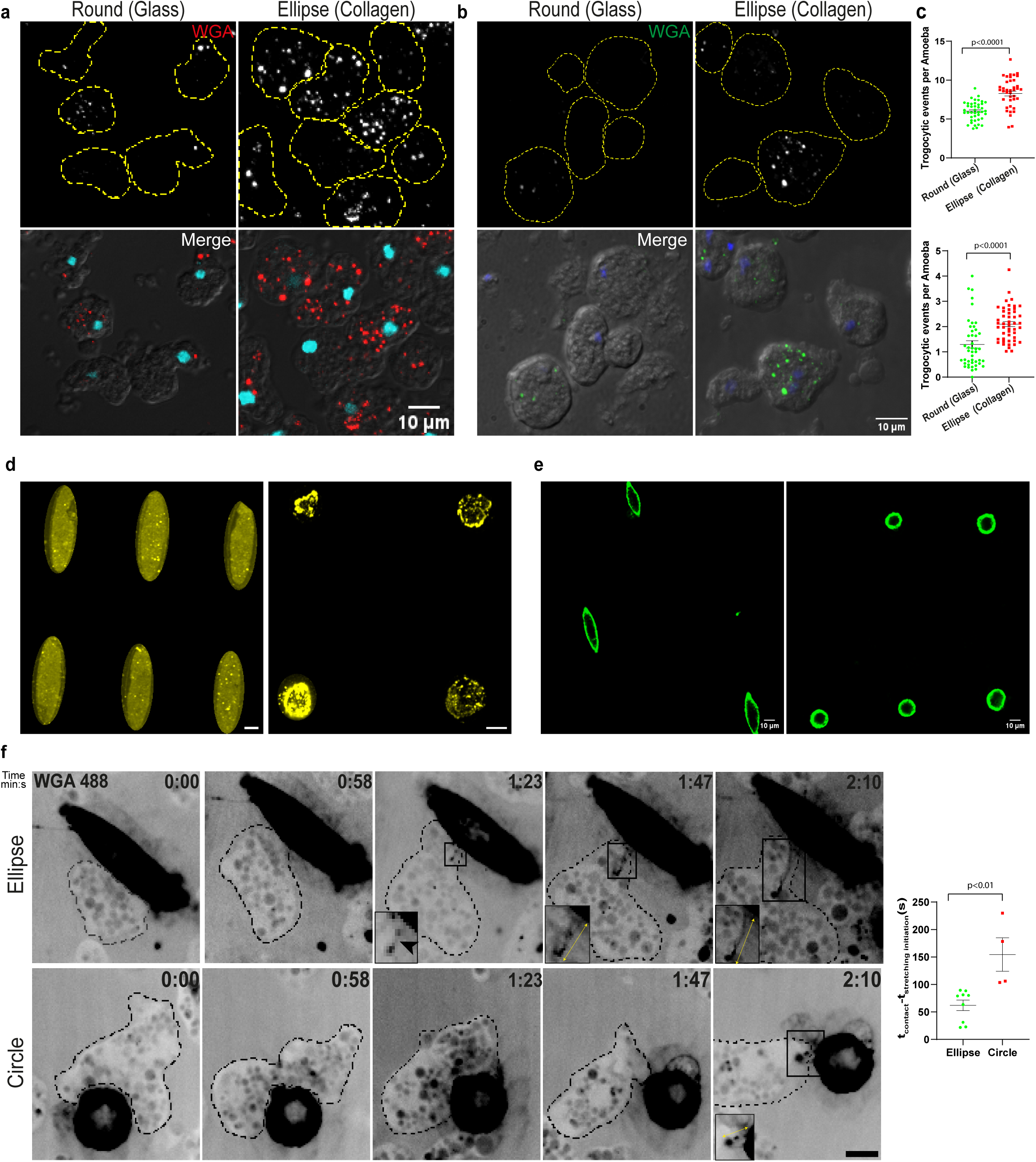
Trogocytosis dynamics depend on the target cell shape. (a) Representative trogocytosis images for the assay performed with SW480 cells seeded on collagen-coated and uncoated surfaces (Glass) after 3h of cell seeding. The cells acquire a round morphology on an uncoated surface and an elliptical morphology (Aspect ratio 4-5) on a coated surface. They were then stained with WGA and Hoechst, incubated with amoeba for 20 minutes, fixed, and analyzed by confocal microscopy. The images show the WGA channel (Red) and the merge channel with DIC. MIPs are shown. Scale bar 10µm. (b) Representative confocal images of trogocytosis with DLD1 cells seeded on uncoated and collagen-coated culture dishes are shown. The Trogocytosis assay was performed after 3 h of cell attachment. Cells were stained with WGA and Hoechst, incubated with amoeba, fixed, and subsequently imaged. Shown are the MIPs with the WGA channel and the Merge channel with DIC. Scale bar 10µm. (Right) Quantification of trogocytosis efficiency is presented for DLD1 cells on glass (round morphology) and collagen (elongated morphology; ellipse AR=4-5). (c) Quantification of Trogocytic efficiency as Trogocytic events per amoeba for SW480 (top) on Glass (Round) and Collagen (Ellipse AR 5). Each data point corresponds to a value from a single field of view. Data from 3 independent experiments were pooled (n = 1404 amoeba), and for DLD1 (bottom). Data were obtained from N = 3 independent experiments, n = 910 amoeba. The mean value and SEM are indicated with black bars. The p-value was derived from an unpaired t-test. (d) Collagen micropatterns of two shapes, Ellipse (left) and Circle (right), were imprinted using PDMS stamps on glass-bottom dishes and visualized using WGA. Scale bar 10µm. (e) Cell morphology after 6h of culture on micropatterned surfaces. Cells were visualized using WGA 488 in spinning disk confocal microscopy. Scale bar 10µm. (f) Trogocytosis time-lapse montages for SW480 cells confined to an elliptical shape and a circle. The WGA channel shows an amoeba undergoing trogocytosis, marked in black outline. The inset shows a magnified view of the region indicated by the box. The black arrowhead marks the stretched membrane tube. The yellow double-headed arrow corresponds to the length extended by the amoeba. Time is displayed as min: sec in the top-right corner of the fluorescent images. The time of contact with the host cell is defined as t = 0. Frames were captured every 1-1.5s. Scale bar = 10µm. (Right) Quantification of the time taken to initiate membrane tubulation for both targets by the amoeba. Each data point represents a value obtained from a single Trogocytic event, n = 9 trogocytic events for the Elliptical target cell (AR 4) and n=4 for the Circle (diameter = 20µm) target cell.

## Discussion

During infection, *E. histolytica* primarily interacts with large, adherent epithelial cells within the intestinal epithelium^46^, where trogocytosis has been implicated in tissue invasion^12^. Indeed, amoebic trogocytosis has primarily been observed in interactions with adherent cell types, such as in the *ex vivo* intestinal tissue^12^ and liver sinusoidal endothelial cells^47^. In the present study, we investigated trogocytosis of adherent target cells, focusing on how target-cell biophysical characteristics influence amoebic trogocytosis. By developing a physiologically relevant monolayer-based trogocytosis assay, we revealed a distinct mechanism of Trogocytic uptake for adherent target cells. Further, we studied the roles of viscoelastic properties and the shape of the target cell in amoebic trogocytosis. To date, only a few studies have examined the influence of target-cell physical properties on trogocytosis vs. phagocytosis primarily in the context of macrophages^15,16^. Of note, recently, Fletcher et al. reported a negative correlation between macrophage-mediated trogocytosis and target-cell cortical stiffness using suspension cells as targets^16^. This mechanical setting, where target cells are presented in suspension, is physiologically relevant for macrophages, whose targets are often suspended immune cells. Thus, the current study reports the first known attempt to investigate how the mechanical properties of host cells could contribute to amoebic trogocytosis using the most appropriate physiological setup - an adherent monolayer.

We uncovered a novel mode of trogocytosis operative in adherent cells, characterized by the formation, elongation, and ultimately scission of membrane tubes from the host-cell surface upon amoebic attachment (Fig. 1). Very recently, a similar cellular process was observed in the interaction between the target cell and the bone marrow-derived macrophage^48^. This is in contrast to the suction-driven mechanism reported for suspension target cells, in which the host cell was stretched into a tunnel-like structure within the trophozoite, and intracellular material was subsequently severed from this region as discrete bites^12,49,50^. After identifying the physical events that constitute trogocytosis of adherent cells, we demonstrated that the shape and viscoelastic properties of target cells affect trogocytosis dynamics. Prior studies have highlighted the role of target cell shape in influencing phagocytosis using artificial substrates such as beads and polystyrene particles^51,52^, but nothing is known in the context of trogocytosis. Here, using micropatterning, we demonstrated the role of target cell shape in trogocytosis using adherent living cells as targets. Real-time imaging data revealed a longer time interval between host-cell contact and membrane-tube initiation for the circular subtype, whereas membrane tubulation in the elliptical target cells occurs preferentially along the major axis (Fig. 4). We propose that this difference arose from the underlying curvature landscape of the target membrane^53^. In circular cells, isotropic high curvature may impose a greater energetic barrier for tube nucleation, compared to the lower curvature along the major axis of elliptical cells.

We therefore propose that cell shape is one of the factors regulating the initiation kinetics of trogocytosis. Given that cell shape and cellular stiffness are often tightly coupled, further studies will be required to delineate the possible contribution of mechanical factors to this process. Trogocytosis is initiated by ligand-receptor interactions between participating cells. This suggests that ligand organization and density can influence the kinetics of trogocytosis initiation. In the context of immune cells, ligand clustering was observed during trogocytosis^54,55^. Our live-cell imaging data analysis revealed a prolonged lag time before pulling, particularly for thicker protrusions. Based on this observation, we hypothesize that this delay may reflect increased time spent on employing an appropriate number of surface ligands at the contact interface.

Post-tube formation, membrane extension dynamics are likely to become increasingly dependent on the mechanical properties of the target cell, including its ability to deform and dissipate stress. Indeed, by integrating live-cell time-lapsed microscopic measurements of membrane extension with a theoretical framework, we found that trogocytosis-induced membrane deformation follows a fiber-bundle like model where each elements behave like a Kelvin–Voigt unit, reflecting that the mechanically pulled target cell membrane exhibits viscoelastic behavior. These findings suggest that altering the viscoelastic properties of the cell might influence trogocytosis dynamics. Given the central role of the actin cytoskeleton in regulating cellular viscoelastic properties, remodeling of actin architecture is therefore likely to modulate the mechanical state of the cell^24–27^. To investigate the effect of cell stiffness, we used multiple actin modulators previously shown to alter cell mechanics^56–59^. Treatment of cells with CK869, myosin inhibition, or depletion disrupted cortical actin and stress fibers^29,33,60^; these effects were consistent with previous reports. These alterations were accompanied by enhanced trogocytosis, whereas α-actinin-4 knockdown, associated with increased stress fiber abundance^40,41^, resulted in reduced trogocytosis. Collectively, these results suggest that trogocytosis increases with a decrease in the stiffness of the cell and are consistent with previous findings^16^. However, Latrunculin A treatment, which produced extremely soft, fluid-like cells, reduced trogocytosis. Further insights gained from the theoretical model predicted that highly fluid-like cells exhibit reduced membrane-stretching dynamics and prolonged scission times, thereby impairing efficient trogocytic uptake. Together, these observations led us to conclude that the relationship between target-cell mechanics and trogocytosis is not simply monotonic. Our interpretation was further supported by results from the trogocytosis assay with cells cultured on substrates matching the stiffness of matrices^42,43^, where increased trogocytosis was observed with cells on an intermediate stiff substrate (100kPa) compared to soft (1kPa) and hard (1000kPa) substrates (Fig. 2- 3). Overall, these findings led us to conclude that Trogocytosis increases with a decrease in the stiffness of the cell up to a certain extent, beyond which it declines, and is different from the recent findings^16,17^. This could be due to distinct regulatory mechanisms operating for target cells in suspension.

However, the increased trogocytosis observed following Jasplakinolide treatment appeared inconsistent with this interpretation, as these cells exhibited fluid-like behavior comparable to that of Latrunculin A-treated cells while largely retaining elastic resistance (E_0_). One possible explanation is that, unlike Latrunculin A treatment, which caused relatively homogeneous disruption of the cortical actin network, Jasplakinolide treatment produced a heterogeneous actin architecture characterized by localized actin aggregates interspersed with regions containing a thinned, but not completely disrupted, cortex (Fig. 2). These locally weakened cortical regions may remain within the optimal mechanical regime for trogocytosis.

Further, our study also revealed that scission preferentially occurs at regions with reduced actin intensity. In most cases, scission was not observed in regions with minimal actin intensity, which may represent highly fluid-like domains within the stretched membrane tube, nor at the enriched actin regions, which may resist rupture (Supplementary Fig. 1). These observations further support the concept that optimal viscoelastic balance regulates trogocytosis not only at the whole-cell level but also locally within the stretched membrane tube. In addition, previous studies have highlighted the role of actin in hindering lipid mobility^61^, and in other related studies, the hindrance was shown to drive friction-mediated scission^62,63^. Further studies will be required to elucidate the molecular mechanism underlying scission. Moreover, our theory-based modeling suggests a conserved mechanism of membrane-tube rupture across distinct mechanical subtypes, in which the breaking rate increases with membrane extension, progressively reducing the tube’s survival probability until rupture occurs beyond a critical threshold.

Together, our study suggests that the mechanical state of the cell regulates multiple stages of trogocytic uptake, including membrane-tube initiation, extension, and scission (Fig. 6), and supports a model in which the mechanical state of the host cell dictates both the mode and the efficiency of ingestion, with amoeba transitioning from phagocytosis of rigid targets to suction-driven or membrane-tubulation–mediated trogocytosis, which relates non-monotonically with cell mechanics. Recently, the KERP2 effector was shown to hijack the host actin cytoskeleton^64^, and previous studies have similarly reported actin cytoskeleton remodeling in epithelial cells on interaction with amoeba ^65,66^. Based on our results, we suggest that the remodeling may help establish a permissive state for productive trogocytosis. Considering the fundamental role of amoebic trogocytosis in augmenting pathogenesis, our results suggest that modulation of host-cell mechanics may represent a potential strategy to influence amoebic virulence.

**Figure 6.**
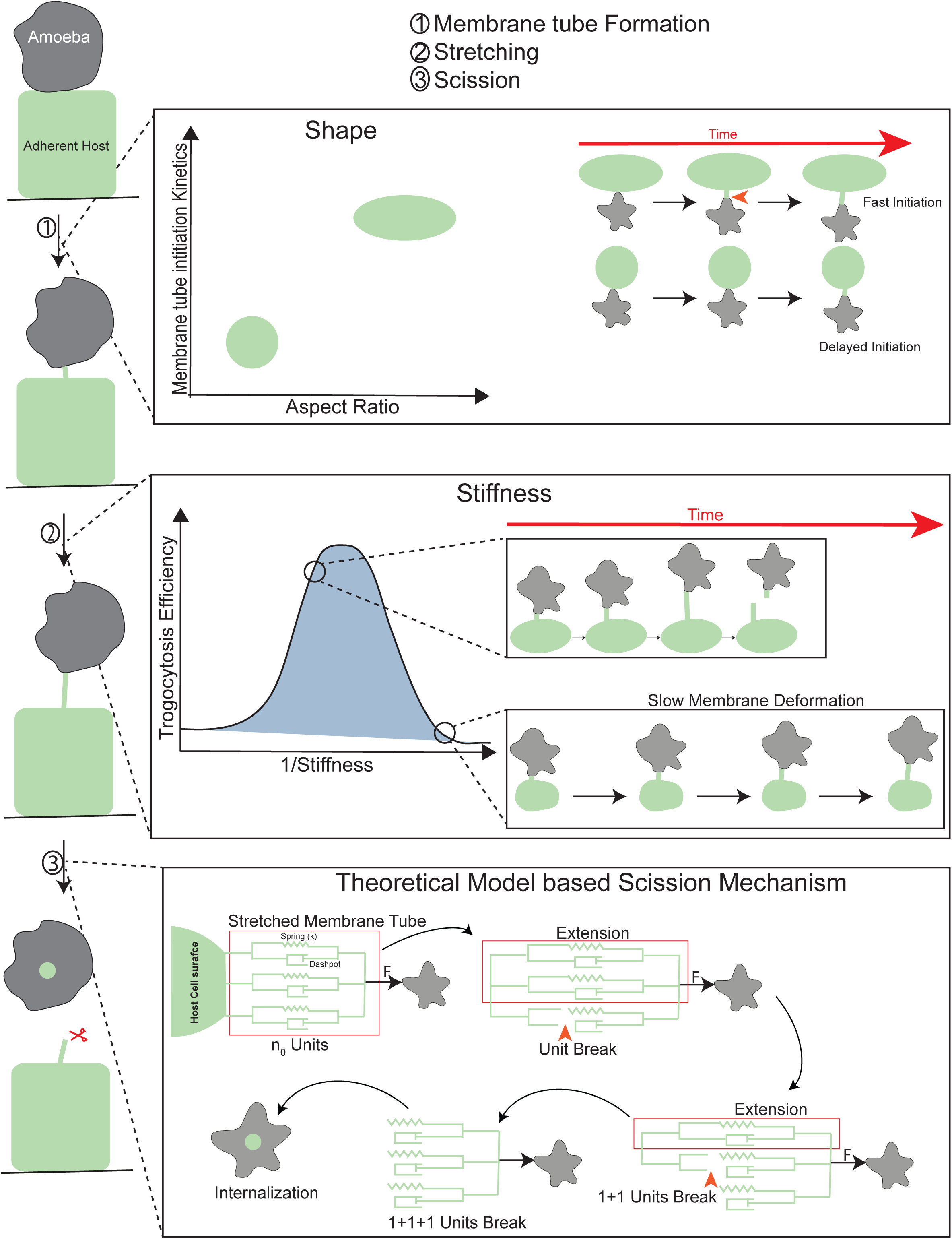
Regulation of Trogocytosis at distinct stages by adherent cell shape and stiffness, together with the scission model. Experimental measurements and analyses indicate that trogocytosis proceeds through membrane tubulation following contact between the amoeba and the target cell. Target cell geometry regulates the initiation kinetics (top), with elliptical cells exhibiting faster initiation dynamics. During tube extension, the deformation rate strongly depends on the mechanical properties of the target cell (middle) and is markedly reduced in highly fluid-like cells. Consequently, trogocytosis efficiency increases with decreasing cell stiffness but declines in extremely fluid-like cells. During the final stage of trogocytosis, tube scission occurs cooperatively, consistent with the theoretical framework (bottom). The spring–dashpot units within the red box represent the elemental units subjected to the applied force (F). As individual units break, the force is redistributed among the remaining units.

## Methods

### Cell culture

1. *E*. *histolytica* strain HM1:IMSS trophozoites were grown axenically in BI-S-33 medium supplemented with 15% (v/v) heat-inactivated adult bovine serum (RM9981), 2.6% v/v vitamin mix, 100U of penicillin/ml, and 100 μg streptomycin/ml (cat. No: 15140122, Thermofisher Scientific) at 35.5° C.

SW480 (CCL-228) and DLD1 (CCL-221) cell lines were purchased from the cell repository of Animal Tissue Culture Cell line (ATCC), respectively, and authenticated for contamination. SW480 cells were maintained in Dulbecco’s modified Eagle medium (DMEM) (cat no. 11995065, Thermofisher Scientific) with 10% fetal bovine serum (FBS) (cat. no. A5256701, Thermofisher Scientific), 100 μg/ml penicillin, 100 μg/ml streptomycin (cat. No: 15140122, Thermofisher Scientific) at 37°C with 5% CO_2_. DLD1 cells were grown in Roswell Park Memorial Institute 1640 (RPMI 1640) (cat. No: 11875093, Thermofisher Scientific) with 10% FBS and 100 μg/ml penicillin–streptomycin at 37°C with 5% CO_2_. All the experiments were performed with cells up to passage number 20.

### Trogocytosis time Lapse Imaging

SW480 cells or DLD1 cells were seeded on a glass-bottom dish (cat no. 100350, SPL) and maintained in their respective cell culture media. Cells were washed with serum-free media and labeled with 5µg/ml WGA (cat no. W11261, Thermofisher Scientific) in serum-free media for 10 minutes prior to imaging. Cells were then washed with the respective serum-free media. For live-cell actin imaging, SW480 cells were labeled with 1 µM SiR actin (cat no. SC001, Spirochrome) and 10 µM Verampil for 1 h, then with 5 µg/ml WGA for 10 minutes, and then washed with serum-free media.

For live cell imaging with Latrunculin A (cat no. 428026, Sigma)-treated host cells. The DLD1 or SW480 cells were seeded on a glass-bottom dish and then washed with serum-free media and treated with 1µM of Latrunculin A diluted in serum-free media, for 2h. The cells were then washed four times with serum free media to remove residual drug. The cells were then labelled with 5µg/ml WGA. Amoeba cells were extracted and washed with BI incomplete media. For trogocytosis live cell imaging, serum-free media in DLD1 or SW480 cells was replaced with BI media containing amoeba and then imaged in an Olympus IXplore spinSR microscope equipped with a 60x oil objective (1.5NA) and a heated stage. Image frame encompassing the WGA channel for Host cell and bright field for amoeba were recorded at the maximum rate obtainable by the system (0.8–2 s interval/frame).

The maximum length stretched was calculated as the length of the membrane tube prior to scission frame and thickness of the extended membrane tube during the initiation of stretching were calculated using the Fiji measurement tools. To calculate the stretching rate, the ratio of net increase in stretched length and total time, was measured.

To calculate Pearson correlation coefficient between actin and membrane distribution across the membrane tube. The line was drawn across the stretched membrane and intensity profile for WGA and Actin profile were calculated. An average background intensity was also calculated by drawing the line in nearby background. Each intensity value is then subtracted with respective background intensity for both the channels. The correlation value then across these intensity profiles was then calculated.

### Trogocytosis Assay

SW480 or DLD1 cells (0.6-1.2×10^5^) were seeded in a 4-well plate in 500 μL complete media. After seeding, cells were incubated at 37 °C with 5% CO_2_ and allowed to achieve 80% confluence. The cells were washed with serum-free media, stained with 5 µg/ml WGA and 2 µM Hoechst 33342 (cat no. I34406, Thermofisher Scientific) for 10 minutes, and then washed 3 times to remove unbound stain. For cells subjected to pharmacological perturbations (Latrunculin A, Cytochalasin D (cat no. C8273), Jasplakinolide (cat no. 420127), Blebbistatin (cat no. B0560), CK869 (C9124), Sigma), staining was performed after treatment. In the case of siRNA or shRNA-transfected cells, the cells were stained for assay after 70h of transfection. The amoeba cells were harvested and washed with BI incomplete media. The amoeba cells were counted and added to the labeled target cell in suspension at a 1:1.1 ratio in a 4-well plate, and the plate was incubated at 37 °C for 30 minutes unless specified. The suspension is then removed, placed on a glass coverslip, and incubated at 37 °C for 30 minutes to allow cells to adhere. The cells were then fixed with 4%paraformaldehyde at room temperature for 15 minutes. After washing with PBS, the coverslips were mounted on a glass slide using Mowiol. Slides were imaged using a spinning-disk confocal microscope (Olympus IXplore spinSR) with a 60× oil-immersion objective (1.5 NA). Image stacks were captured using 0.75-1 μm *z*-sectioning. To avoid user bias, images were captured randomly using the Hoechst 33342 channel rather than the WGA channel. For analysis, z-stacks of the cell were combined to generate a maximum intensity projection (MIP) using ImageJ software, and the resulting image was analyzed using Motion Tracking software (http://motiontracking.mpi-cbg.de/).

The WGA labeled objects were identified as vesicles depending on their size and intensity in each frame. The objects within the amoeba were considered for statistics by masking the amoeba and the objects colocalizing with Hoechst channel was not considered as trogocytosis events and were omitted. The total number of WGA objects were then divided by total number of amoeba in the frame to get Trogocytic events per amoeba, which is used as the measurement of trogocytosis efficiency.

### Atomic Force Microscopy to measure cell stiffness

#### Indentation Experiments

The indentation experiments were performed on single cells, using a rigid spherical indenter, using an atomic force microscope (JPK Nanowizard II, Berlin, Germany) to determine their viscoelastic properties. To avail the spherical indenter, commercial tipless rectangular cantilever (Micromash, Bulgaria) was customised by attaching a 5μm(diameter) glass bead using the liftoff method [10]. Cantilevers with a typical force constant of 0.3Nm−1 were used. The force constant of the cantilever(kc) was determined before each experiment using the thermal calibration method provided in JPK Nanowizard II. Cells were seeded on coverslips and maintained in their culture condition for 24h. AFM studies were performed with wild-type cells and cells treated with different pharmacological interventions (10 µM Cytochalasin D, 1 µM Latrunculin A, 0.2 µM Jasplakinolide, and 75 µM Blebbistatin). Cells treated with DMSO served as the control. The cells were then washed 3 times to remove residual drug prior to AFM, to mimic the conditions used for the trogocytosis assay. The approach–retract experiments were performed on single cells at an extension speed of 2 µms-1 over a 3 × 3 µm2 area using a 6 × 6 grid, with a sampling rate of 2 kHz. The high-quality curves, characterized by a smooth approach and retract profile with minimal drift, were analyzed. The mean values for each rheological parameter were then obtained and compared using an unpaired t-test or One-way Anova to determine statistical significance. The indentation was kept small (≈ 700 nm). The small indentation on the cells is important to avoid the bottom-surface effect, as the cells have a finite thickness [5, 1]. The typical height of a SW480 and DLD1 cell line is 8-10 µm. The experimentally recorded parameters (raw Data) are cantilever deflection (dcant, in volts) and the base-piezo displacement (dpz, in volts). dcant and dpz were converted into units of meters by multiplying them by photodetector sensitivity and base-piezo sensitivity, respectively. The force (F) on the cantilever/sample was calculated by multiplying dcant by the cantilever Force constant. The deformation of cells was determined by subtracting the base piezo displacement from the cantilever deflection (δ = dpz − dcant).

### Model Used

Ting’s model provided an analytical solution for linear viscoelastic contact between a half-space and an axisymmetric indenter, applicable to any loading history ^22,23^. This is relevant to conventional AFM experiments, in which cells are deformed at a constant loading rate. Efremov et al. developed a complete analysis protocol to determine the viscoelasticity of a single cell by fitting Ting’s model on conventional AFM force–indentation curves^67^. We used a MATLAB code developed by Efremov et al. ^23^ to fit the force curves using Ting’s model and assess the cell viscoelasticity. The code is available at https://doi.org/10.5281/zenodo.10529399. The entire curve, encompassing both the approach and retract phases, was analyzed using Ting’s model. We used constitutive equations that account for time-dependent Young’s modulus, specifically power-law rheology (PLR). The relevant parameters in these equations were determined to provide a comprehensive description of the cell’s viscoelastic behavior. In Ting’s description, the relation between force F (t, *δ*(*t*)) and indentation on the sample *δ*(*t*) is given by

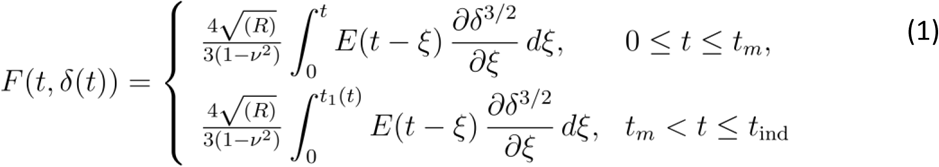

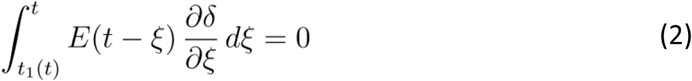

where R and *v* are the radius of the indenter and Poisson’s ratio of the sample, respectively. t is the time initiated at first contact, *t*_*m*_ is the time duration of the approach part and *t*_*ind*_ is the time of the full approach-retract cycle while the tip is in contact with the substrate. *ξ* is the dummy time variable used for the integration and *t*_1_(*t*) is the auxiliary time that is determined from eqn (2). E(t) is the relaxation modulus and can be used from any linear viscoelastic constitutive equations that describe the sample behaviour. For fitting, the Poisson ratio (*v*) was taken as 0.5 for each cell.

The constitutive equation with power law relaxation (PLR) describing the mechanics of soft glasses is

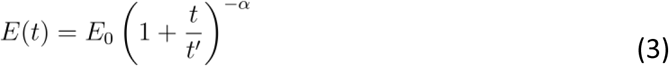

where *E*_0_ and *α* are the instantaneous Young’s modulus and power-law exponent, respectively. *t*^′^ is a small-time offset and is set to the sampling time (inverse of the sampling rate, which was 2 kHz for all experiments). The value of *t*^′^ used in our analysis is 5 × 10^−4^ s.

Equation (3) has been modified to account for the long-term non-zero relaxation modulus *E*_∞_. The relaxation modulus for the modified power law rheology (mPLR) model is given by

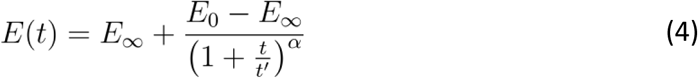

We used Equation (3) (simple PLR) for the data analysis as we found *E*_∞_ ∼ 0 for the cells under investigation. The use of Equation (3) reduces the program run time during the analysis. The Power law exponent (α) and E_0_ values were determined for all the conditions.

### Actin Staining

For imaging cytoskeletal actin to verify the effect of pharmacological interventions on actin. SW480 or DLD1 cells were seeded on glass conditions and were maintained in their culture conditions for 48-72h after which cells were washed with respective serum free media and different pharmacological perturbations were carried by treating cells with actin modulating drugs (1µM Latrunculin A, 0.2µM Jasplakinolide, 10µM Cytochalasin D, 40µM CK869 and 75µM Blebbistatin), diluted in serum free media and incubating for respective time. Cells treated with DMSO were used as a control. For washout the cells were washed four times with serum free media after treatment and further kept in serum free media for 1 hour in the absence of drug. For F actin visualisation in siRNA and shRNA transfection, cells were fixed after 70h of transfection. The cells were then fixed with 4% paraformaldehyde at room temperature for 15min, followed by three washes with 1X PBS. Cells were then permeabilized with 0.1%Triton for 10mins. Samples were then stained with Phalloidin AF568 (cat no. A12380, Thermo Fisher Scientific or AF647 (cat no. 82287, Thermo Fisher Scientific) (1:100 dilution) for 1h. Following staining sample were washed thrice with 1X PBS. Coverslips were then mounted on glass slides using Mowoil and imaged using FV3000 Olympus microscope at 63x oil immersion objective lens.

### siRNA and shRNA transfection

OntargetPlus siRNA targeting the NMIIA (information included in the Supplementary file Table I) was purchased from Dharmacon. The siRNA was used at a working concentration of 25 nM. Cells were seeded 24 h before performing siRNA transfection. The manufacturer’s protocol was used for transfection using Dharmafect (cat no. T-2001-03, Dharmacon) as the transfection reagent.

Alpha actinin 4 shRNA clone (information included in Supplementary file Table II) was part of the MISSION shRNA product line from Sigma-Aldrich. The TRC1.5 pLKO.1-puro non-mammalian shRNA Control Plasmid DNA (SHC002) is a negative control containing a sequence that should not target any known mammalian genes but engage with RISC. Cells were seeded 24 h before performing shRNA transfection. Cells were transfected at 50–60% confluency with 0.5 µg of shRNA plasmid DNA using Lipofectamine 3000 (cat no. L3000008, Thermo Fisher Scientific) transfection reagent.

### Protein Preparation and Western Blotting

Cells were collected and lysed using a buffer containing 50 mM Tris-HCl (pH 7.4), 150 mM NaCl, 2 mM DTT, 1% NP40, and 1 mM EDTA, supplemented with 10 µg/ml protease inhibitor cocktail (PIC, Sigma). The lysate was clarified by centrifugation at 19,000 ×g for 15 minutes at 4°C to remove cellular debris. Protein levels were determined using the Bradford assay. Samples were then mixed with 1× SDS loading dye and heated at 95°C. Proteins were resolved by SDS-PAGE and subsequently transferred onto a 0.45 µm nitrocellulose membrane (GE Healthcare, Cat. 10600002). The membrane was blocked with 5% BSA and incubated with the primary antibodies (rabbit anti-NMIIA at 1:800 dilution (cat no. M8064, Sigma), Mouse anti-Tubulin at 1:850 dilution (cat no. T6557, Sigma) for 2 hours at room temperature. Following this, fluorescently labeled secondary antibodies were applied, and protein bands were visualized using the Li-Cor Odyssey Infrared Scanning System.

### RNA extraction and quantitative real-time PCR

Total RNA was extracted from the cells using the RNAeasy kit (cat no. 74104, QIAGEN), and cDNA was prepared using the High-Capacity RNA-to-cDNA kit (Life Technologies, Cat# 4387406). Real-time qPCR reactions (sequence information in Supplementary file) were performed using the SYBR Green Kit (cat no. 4367659, Thermo Fisher Scientific) and corresponding primers on Applied Biosystems 7300 Real-Time PCR System or Thermo Quant Studio 3.0.

### Polyacrylamide Substrates

Preparation of Polyacrylamide substrates was carried out following the previously reported protocols ^68,69^. Briefly, glass coverslips were rendered reactive by sequential treatment with NaOH and 3-aminopropyltrimethoxysilane (APTMS) (cat no. 281778, Sigma), followed by glutaraldehyde treatment to enable covalent attachment of the PA gels to the glass surface. PA gels were prepared using acrylamide at final concentrations of 2.5, 12, and 22 wt/vol%, together with corresponding bis-acrylamide concentrations of 0.08, 0.4, and 0.75 wt/vol%, respectively, to generate soft (1kPa), intermediate-stiffness (100kPa), and stiff substrates (1000kPa). Polymerization was initiated by the addition of ammonium persulfate (APS; 1/100 total volume) and tetramethyl ethylenediamine (TEMED; 1/1000 total volume) to the gel precursor solution. Approximately 80 μl of gel solution was placed at the center of an amino-silanated glass coverslip (12 mm diameter), and a chloro-silanated coverslip (Treated with Sigmacote (cat no. SL2, Sigma) was carefully positioned on top. After 30 minutes of polymerization, the top coverslip was removed. Residual monomers and crosslinkers were eliminated by washing the gels with PBS. For extracellular matrix functionalization, collagen I was coupled to the hydrogel surface using the heterobifunctional crosslinker Sulfo-SANPAH (cat no. A35395, Thermo Fisher Scientific). Briefly, 200 μl of 0.2 mg ml⁻¹ Sulfo-SANPAH prepared in 50 mM HEPES (pH8.5) was added to the gel surface, followed by UV illumination for 30 min inside the Hood. The gels were subsequently washed 3 times with 50 mM HEPES (pH 8.5) in PBS. Activated gels were then coated with 500 μl of 0.05 mg ml⁻¹ rat tail type I collagen (cat no. A10483, Thermo Fisher Scientific) prepared in acetic acid and incubated for 3 h at 37°C. Before cell seeding, the hydrogels were sterilized under UV light in a laminar flow hood for 20 min, then washed with PBS and culture medium. For trogocytosis assays, DLD1 cells were trypsinized and seeded onto the polyacrylamide substrates at a density of 1.7×10^5^. The assay was performed 24 h after cell seeding. To assess cell morphology, cells were fixed and phase-contrast images were acquired using a 10× air objective on a Nikon Eclipse Ti2-E inverted microscope.

### Micropatterning

Collagen micropatterns were generated on glass-bottom dishes. using PDMS stamps. Polydimethylsiloxane (PDMS) stamps were prepared from master molds and stored in a dust-free environment until use. Before collagen printing, the stamps were treated in a UV ozone cleaner (Holmarc) for 45 min. The stamps were inverted onto Collagen solution, and incubated for 1 h at 37°C in a humidified chamber. Collagen solution was used at 50 µg ml⁻¹ in phosphate-buffered saline.

35mm Cell culture Glass-bottom dishes were similarly treated in the UV ozone cleaner for 1 h before printing. The collagen-coated PDMS stamps were immediately brought into conformal contact with the glass bottom dish to transfer the pattern. A small weight was placed on top of each stamp to ensure uniform contact. Following stamping, the stamps were carefully removed, and non-patterned regions were passivated with Pluronic F127 (cat no. P2443, Sigma; 0.4% w/v in PBS) for 30 min. The substrates were then washed three times with PBS. Cells were subsequently seeded at a density of 10,000-15,000 cells ml⁻¹ and incubated for 6 h to allow attachment and spreading. Cells were then prepared for live-cell imaging of trogocytosis.

### Statistical Analysis

Sample sizes and statistical tests are included in the figure legends. All statistical tests were conducted using GraphPad Prism. The results are expressed as either mean ± standard deviation (SD) or mean ± standard error of the mean (SEM). Significance was tested compared to the control groups and defined as significant when *p* ≤ 0.05 or non-significant when *p* > 0.05.

## Author Contributions

T.A. and S.D. designed the project. B.B. and S.K.N. performed data analysis and theoretical modeling. T.A., Shu.P., and Shi.P. performed AFM measurements and analysis. K.R. performed knockdown experiments. T.A., N.P., N.G., A.D., and S.T. performed microfabrication experiments. T.A., S.D., B.B., S.K.N., Shu.P., Shi.P., N.P., and N.G. contributed to writing and editing the manuscript and figures.

## Acknowledgements

We thank our laboratory technician, Ms. Rabiya Naaz, for continuous technical assistance. We also thank Dr. Rati Sharma for providing access to the UV ozone cleaner and Dr. Shamik Sen for valuable experimental suggestions. We gratefully acknowledge Dr. Yannis Kalaidzidis for assistance with the motion tracking analysis and Dr. Peter Sollich for insightful discussion. This work was supported by the Department of Biotechnology (DBT) grant BTPR50294COT142662023 and a University Grants Commission Fellowship. We acknowledge the DST-supported FIST facility at IISER Bhopal for infrastructure support for live-cell imaging experiments. We also acknowledge the Central Instrumentation Facility at IISER Bhopal for access to the confocal microscopy facility.

