## Supplementary File for "Cell Mechanics Regulate Membrane Tubulation-Driven Trogocytosis in Adherent Cells"

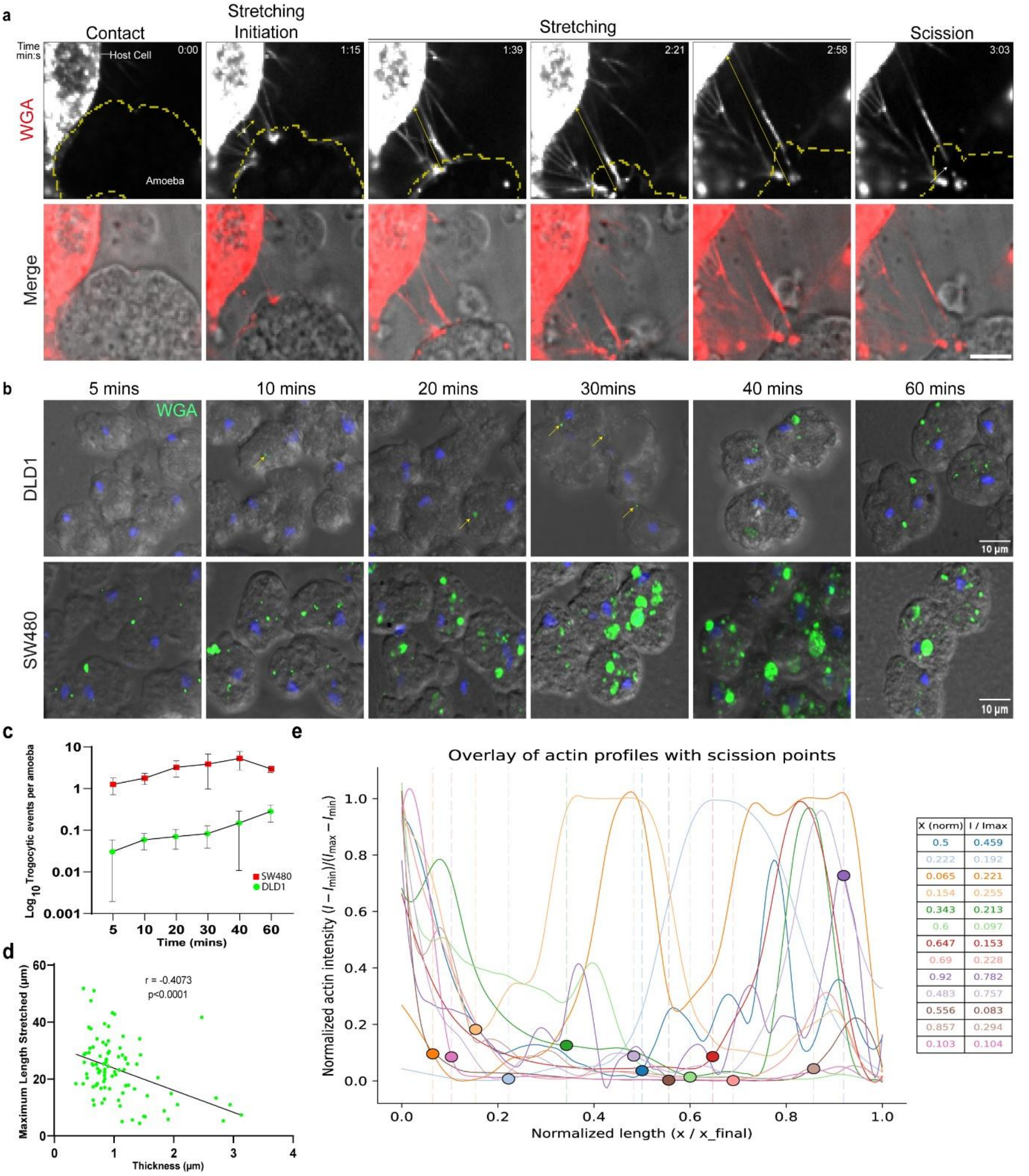


**Supplementary Figure1. Comparative profiling of trogocytosis.** (a) Representative time-lapse montages show amoebae undergoing trogocytosis of live DLD1 cells. WGA-labeled DLD1 cells were incubated with amoeba and imaged using spinning-disk microscopy, with frames acquired every 1s. The trogocytic event is indicated by a yellow arrow, while the double-headed yellow arrow corresponds to the extension of protrusion by the amoeba. The WGA fluorescence channel and its overlay with the bright-field image are shown. The amoeba engaged in trogocytosis is outlined in yellow, and scission is marked by a white arrow. Time is displayed in min:s in the top right of the fluorescence images, with initial contact defined as t = 0. Scale bar: 10 µm. (b-c) Time course of Amoebic Trogocytosis with SW480 and DLD1 host cells. Cell monolayers were labeled with the membrane stain WGA and the nuclear stain Hoechst. These labeled monolayer cells were then incubated with amoeba for the indicated time point at 37 $^{oC}$. Representative trogocytosis images at different time points are shown. An overlay of WGA (Green) and DIC is presented. Scale bar represents 10µm (b) and Quantification of trogocytosis efficiency over time. Data are presented as mean ± SEM from a single experiment, measured across six time points (c). (d) Plot of extended tube thickness versus maximum extension length. Each data point represents a value obtained from a single live trogocytic event, with a total of n = 91 events pooled from SW480 and DLD1 cells. The Pearson correlation coefficient (R) and p values are indicated. (e) Normalized actin intensity profiles along membrane tubes are shown for multiple datasets, with position along the tube scaled to the final tube length. Actin intensity for each dataset was normalized using min–max scaling $(I-I_{min})/(I_{max}-I_{min})$, enabling comparison across tubes. Profiles were interpolated onto a common spatial grid and smoothed using a Savitzky–Golay filter for visualization. Scission positions, identified from experimental annotations, are indicated by filled circles and corresponding vertical dashed lines. The table on the right summarizes the normalized position (X) and actin intensity (I) values at the scission point for each dataset, with colors matching the respective curves. A total of 13 time-lapse videos of trogocytic events were analyzed.

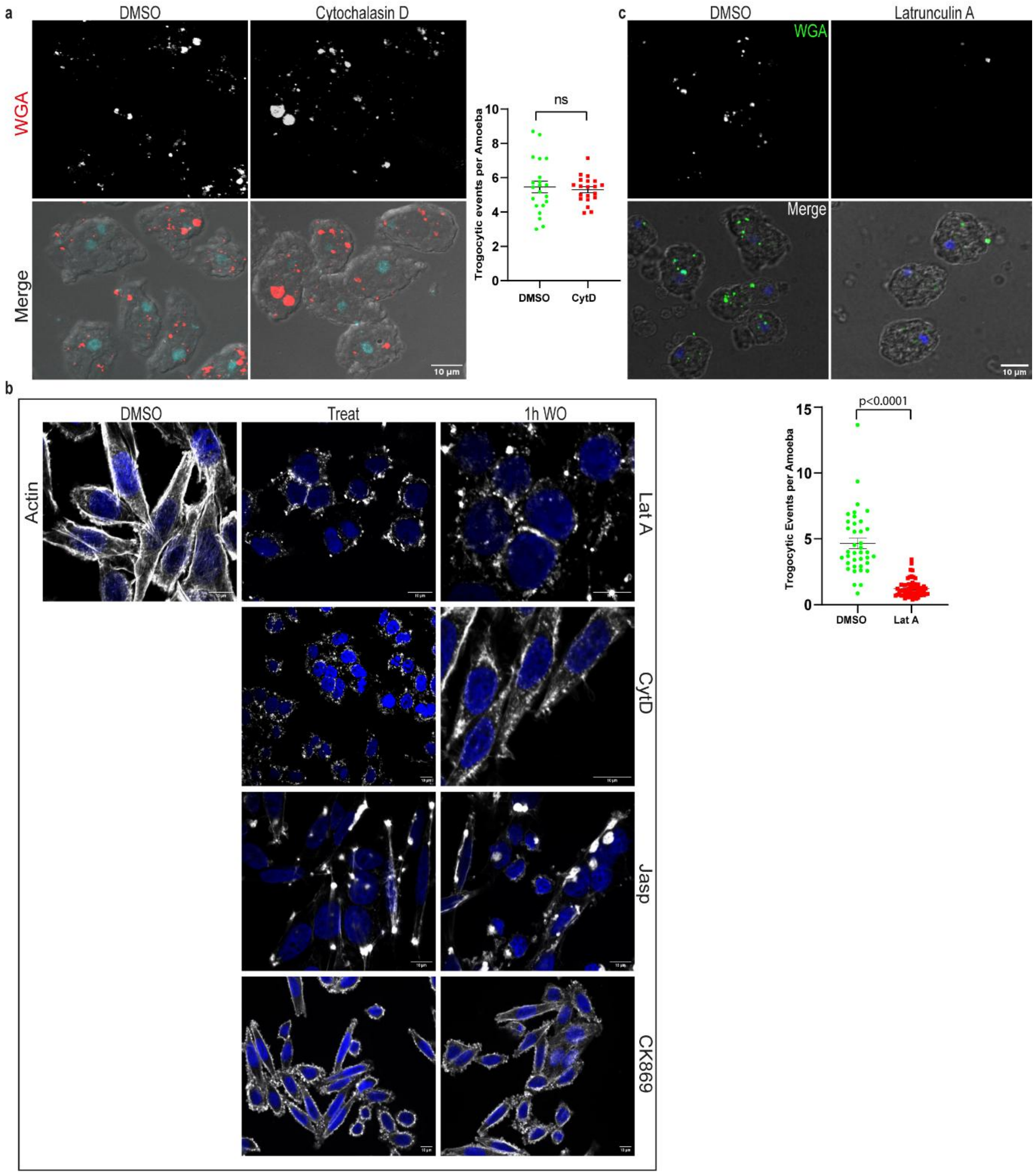


**Supplementary Figure2. Effect of actin modulating agents on Actin architecture and Trogocytosis** (a) The left panel shows the representative confocal images of amoebic trogocytosis in SW480 cells treated with 10 µM Cytochalasin D or DMSO for 4 h. Cells were stained with WGA (membrane) and Hoechst (nuclei), incubated with amoebae for 40 min, and fixed. The WGA channel (red) and its merge with DIC are shown. Scale bar, 10 µm. The right panel shows the quantification of trogocytosis efficiency. N=3 independent experiments are pooled together. Each data point corresponds to a single frame, n≈644 amoeba for both conditions. Mean and SEM are indicated with black lines. The statistical significance was assessed using an unpaired t-test. (b) SW480 cells were treated with DMSO (vehicle) or the indicated actin modulators in serum-free medium. For washout experiments, cells were washed 3–4 times with serum-free medium to remove residual drug and incubated for an additional 1 h. Cells were then fixed, permeabilized, and stained with Phalloidin-568 to visualize F-actin and DAPI to label nuclei. Images were acquired by confocal microscopy. Actin is shown in grayscale, and nuclei (DAPI) in blue. (c) Representative images of trogocytosis (Top). DLD1 cells were treated with either Latrunculin A for 2 hours or DMSO, followed by staining with the membrane marker WGA and the nuclear dye Hoechst. These cells were then incubated with amoebae for 30 minutes before fixation. Imaging was performed using confocal microscopy. The panel displays the WGA channel (green) and its overlay on the DIC image to visualize the amoeba. Scale bar: 10 µm. (Bottom) Microscopy-based quantification of trogocytosis efficiency. Data from three independent experiments (N = 3) were pooled. Each data point corresponds to a single frame, and a total of n = 1265 amoebae were analyzed per condition. The mean and SEM are indicated by black lines. Statistical significance was assessed using an unpaired t-test.

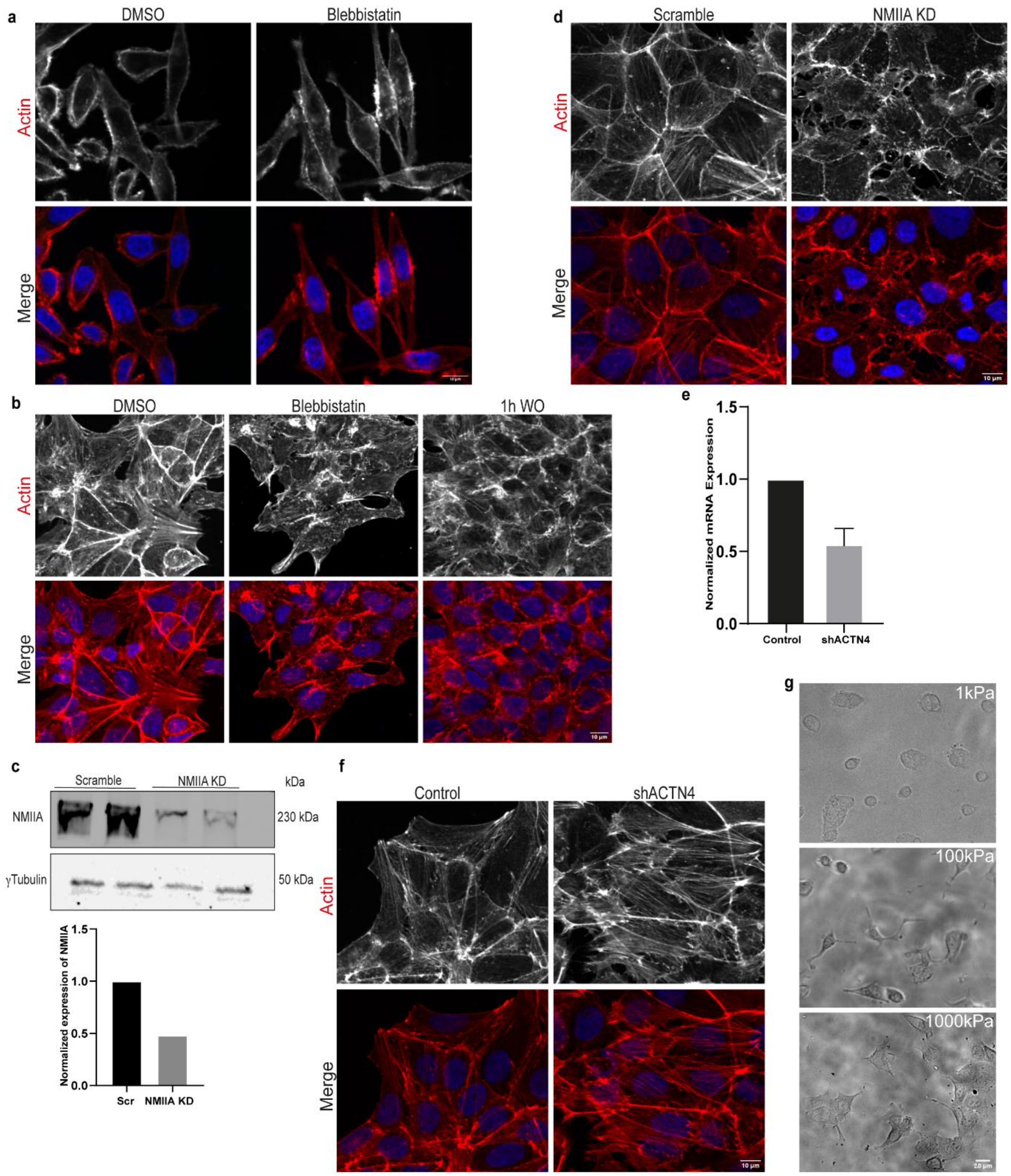


**Supplementary Figure3. Stress Fiber Organization in response to actomyosin modulations** (a) Confocal images show actin staining in SW480 cells treated with the myosin inhibitor Blebbistatin (75 µM) and an equivalent amount of DMSO for 4 h. The cells were then fixed, permeabilized, and stained with Phalloidin 568 to visualize actin architecture and DAPI. The images show actin staining in gray scale and in merged (Red) with DAPI (Blue). Scale bar 10µm (b) Confocal micrographs depict the actin cytoskeleton in DLD1 cells following treatment with the myosin inhibitor Blebbistatin (75 µM) or DMSO control for 4 h. After treatment, cells were fixed, permeabilized, and labeled with Phalloidin 568 to visualize filamentous actin, along with DAPI for nuclear staining. For washout experiments, cells were washed three times with serum-free medium to remove residual drug, then incubated for an additional 1 h in fresh, drug-free medium. Actin is shown in grayscale and in red in the merged image, and nuclei are shown in blue. Scale bar: 10 µm. (c) (Top) Validation of knockdown efficiency in DLD1 cells transfected with NMIIA-targeting siRNA or a scrambled control was performed by Western blotting. Cells were harvested 70 h post-transfection for analysis, with γ-tubulin used as a loading control.

(Bottom) Quantitative analysis of band intensities is shown to assess knockdown efficiency, with NMIIA levels normalized to the tubulin loading control (N=1). (d) Fixed-cell images of Scramble and NMIIA-depleted DLD1 cells stained for actin filaments (Phalloidin AF 568). (e) ACTN4 expression was silenced in DLD-1 cells using ACTN4-targeting shRNA, with a non-targeting shRNA as a control. Knockdown efficiency was validated by quantitative reverse transcription analysis, with ACTN4 levels normalized to GAPDH. Data represent the mean ± SEM from three independent experiments. (f) Stress fiber organization visualized by F- actin staining in DLD1 cells transfected with control shRNA and ACTN4 shRNA. Scale bar: 10 µm (g) Representative phase contrast images of DLD1 cells cultured on 1kPa, 100 kPa, and 1000kPa polyacrylamide gels after 24 hours of cell seeding.

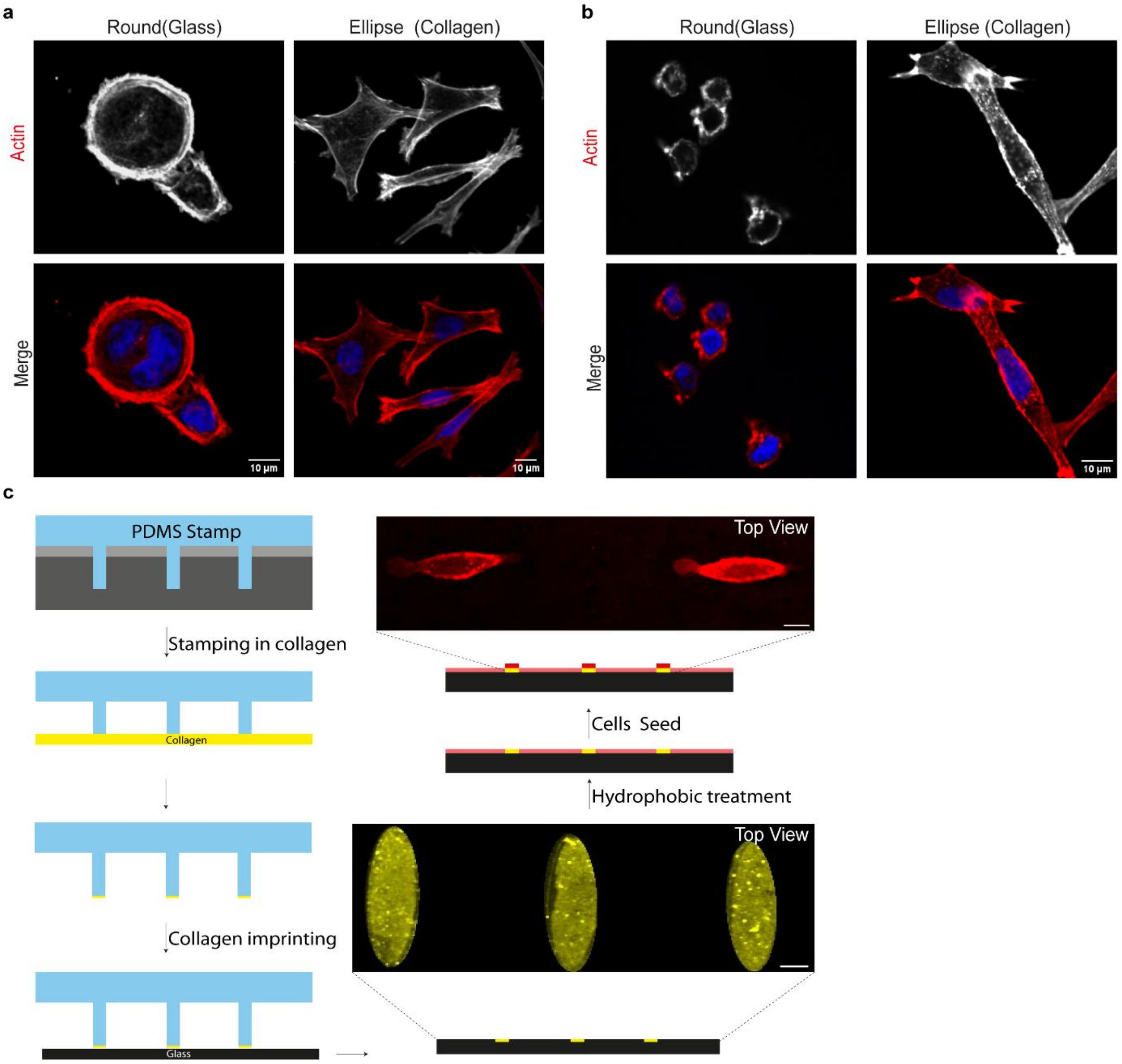


**Supplementary Figure4. Actin organization on collagen substrate and micropatterning** (a-b) Representative actin staining images for SW480 (a) and DLD1 (b) cells seeded on collagen-coated and uncoated (glass) surfaces. Cells adopt a rounded morphology on uncoated surfaces and an elongated, elliptical shape (aspect ratio ~4–5) on collagen-coated substrates. Cells were fixed after 3h of seeding, permeabilized, stained with Phalloidin 568 to visualize filamentous actin, DAPI for nuclear labeling, and imaged by confocal microscopy. The actin channel is shown (red) and DAPI in blue. Scale bar: 10 µm. (Right). (c) Microfabrication method scheme. PDMS stamps (Blue) were fabricated by replica molding from silicon masters (Grey). Prior to microcontact printing, stamps were treated with UV ozone cleaner for 40 minutes. Stamps were then incubated with rat tail collagen (50 µg $ml^{-1}$ in PBS) (yellow) for 1 h, and brought into conformal contact with glass-bottom dishes (black). The magnified image showed an imprinted collagen pattern on the surface of glass-bottom dishes. Subsequently, non-patterned regions were passivated with 0.4% (w/v) Pluronic F127 in PBS for 30minutes (light red). For cell micropattern generation, cells (red) were then trypsinized and seeded onto the imprinted collagen patterns. A magnified view shows SW480 cells spread on elliptical micropatterns and stained with WGA for visualization. Scale bar 10µm.

**Supplementary Table**

**Table 1: Detailed information about siRNA and shRNA used in Figure 3. For knock down**

| Gene Name | **Gene ID** | **Reagent Type** | **Sequence** | **Catalogue Number** | **Company** |
|---|---|---|---|---|---|
| MYH9 | 4627 | ON-TARGET plus siRNA | GUAUCAAUGUGACCGAUUU | J-007668-05-0002 | Dharmacon |
| ACTN4 | 81 | shRNA | CCGGCCCGTATAAGAACGTCAATGTCTCGAGACATTGACGTTCTTATACGGGTTTTTG | TRCN0000055786 | Sigma |

**Table 2: Primers used for qPCR to check shRNA knockdown efficiency**

| **Gene Symbol** | **Forward Primer** | **Reverse Primer** |
|---|---|---|
| **ACTN 4** | TTCAACCACTTCGACAAGGA | ATGAGGCAGGCCTTGAACT |

**Supplementary Video Legends**

**Supplementary Video 1. Live-cell imaging of trogocytosis in SW480 cells.**
Time-lapse microscopy showing amoebic trogocytosis of SW480 cells stained with WGA AF488 (green). The video displays a merged view (Right) of bright-field images of the amoeba and the WGA signal, while the WGA channel alone is additionally shown in grayscale (Left).

**Supplementary Video 2. Live-cell imaging of trogocytosis in DLD1 cells.**
Time-lapse microscopy showing amoebic trogocytosis of DLD1 cells stained with WGA AF594 (red). The video displays a merged view (Right) of bright-field images of the amoeba and the WGA signal, while the WGA channel alone is additionally shown in grayscale (Left).White arrowhead marks the trogocytic event. DLD1 cells exhibit prolonged membrane stretching during trogocytosis.

**Supplementary Video 3. Recoil of the membrane tube following scission.**
Time-lapse microscopy showing recoil of the extended membrane tube back toward the SW480 cell surface after scission. SW480 cells were stained with WGA AF488 (green). The WGA channel (Left) is shown in grayscale and merged (Right) with bright-field to visualize the amoeba.

**Supplementary Video 4. Retraction of the membrane tube without scission during trogocytosis.**
Time-lapse microscopy showing retraction of the stretched membrane tube in the absence of scission, representing an unsuccessful trogocytosis event. SW480 cells were stained with WGA AF488 (green). The WGA channel (Left) is shown in grayscale and merged (Right) with bright-field to visualize the amoeba.

**Supplementary Video 5. Actin localization within the stretched membrane tube during trogocytosis.**
Time-lapse microscopy of SW480 cells showing the presence of actin within the stretched membrane tube during amoebic trogocytosis. SW480 cells were stained with WGA AF488 (green) and SiR-actin (magenta). The SiR-actin channel is additionally displayed in grayscale (Left).

**Supplementary Video 6. Live-cell imaging of trogocytosis with elliptically constrained SW480 cells.**
Time-lapse microscopy of SW480 cells confined to an elliptical geometry during amoebic trogocytosis. SW480 cells were stained with WGA AF488 (green). The WGA channel is shown in grayscale (Left) and merged (Right) with bright-field to visualize the amoeba.

**Supplementary Video 7. Live-cell imaging of trogocytosis with circularly constrained SW480 cells.**
Time-lapse microscopy of SW480 cells confined to a circular geometry during amoebic trogocytosis. SW480 cells were stained with WGA AF488. The WGA channel is displayed using an inverse LUT. The amoeba is visible owing to autofluorescence.

### Supplementary Notes

#### I. Model:

Motivated by the experimental data, we model the host cell protrusion as a parallel assembly of $n_0$ Kelvin-Voigt (KV) units (Fig 4a). The model falls into the larger class of fiber-bundle models[1,2]. Each KV unit consists of a spring, with a spring constant $k$, and a dashpot, with friction coefficient $\xi$, connected in parallel. Assuming that $\ell(t)$ is the extension of a KV unit from its natural rest length, the force exerted on a single KV unit can be written as

$$f_{\text{unit}}(t) = k\ell(t) + \xi\dot{\ell}(t) \tag{1}$$

where $\xi = 6\pi\eta R$, $\eta$, the fluid viscosity and $R$, the radius for the KV unit. For simplicity, we set $R = 1$ as the unit of length. The mechanical parameters $k$ and $\eta$ will depend on the cellular states and other physical conditions, such as cell shape and interaction with the substrate. The total force on the protrusion is

$$F(t) = n_0 f_{\text{unit}}(t) = n_0[k\ell(t) + \xi\dot{\ell}(t)], \tag{2}$$

where $n_0$ is the total number of KV units that form the protrusion.

For a specific protrusion, we assume that the applied force by amoeba, $F(t)$, remains constant, $F_0$. Solving Eq. (2) yields $\ell(t) = \frac{F_0}{n_0 k}\left(1 - e^{-\frac{kt}{\xi}}\right)$, where we have assumed that the stretching length is zero at $t = 0$. In practice, the 'start' of the process is hard to determine, and we find a non-zero protrusion length when we start our measurements. To account for this, we assume that $\ell(t = 0) = \ell_0$, where $t = 0$ designates the start of the measurement. Then, we can write

$$\ell(t) = a(1 - e^{-bt}) + \ell_0, \tag{3}$$

where $a = F_0/n_0 k$, and $b = k/\xi$. We first checked if the model was consistent with the data. In the experiment, we have measured the time-dependent data of $\ell(t)$. We fit the data with Eq. (3) and obtain the parameters $a$, $b$ and $\ell_0$. Therefore, plotting $(\ell(t) - \ell_0)/a$ versus $bt$ reveals that the data collapse onto a master curve, as shown in Fig. 4(c). This excellent data collapse demonstrates the validity of the basic model. We can now use the model to gain further insight into the process.

#### II. Tube breaking mechanism:

As detailed in the main text, we propose a nucleation-like mechanism for the failure of the protrusion. Given a length $\ell$ of a KV unit, it can break at any point throughout the length. Breaking occurs when the applied force overcomes the binding within the KV unit, which can be viewed as comprising monomeric units, leading to rupture. Therefore, the rate of such a breakage will depend on the applied force: $r(\ell) = \bar{r}_0 \exp[\bar{\alpha} f_{\text{unit}}(\ell)]$, where $\bar{\alpha}$ is related to the binding energy between subsequent monomers and $\bar{r}_0$ is a constant[3,4]. As shown in the last section, $f_{\text{unit}}(\ell) = k\ell + \xi v(\ell)$. The experimental data show that the mean velocity remains constant for the process; therefore, treating $v$ as a constant, we can write $r(\ell) = r_0 \exp[\alpha \ell]$, where $r_0$ and $\alpha$ are two constants. The constant velocity requires that the amoeba apply a larger force to a thicker protrusion than to a thinner protrusion. Within our simple model, one possible mechanism is that the amoeba first employs specific ligand molecules to attach to each of the units in a protrusion before it starts pulling. The pulling force is exerted by these ligands. Therefore, the total force is proportional to the number of KV units in the protrusion.

Since the KV units are arranged in parallel, the probability of their breaking progressively increases as more of these units break. When one of the units breaks, the force is redistributed in the rest. Since an increase in force increases the probability of breaking exponentially, it will be a fast process; we assume that the protrusion breaks as soon as one of the KV elements breaks (weakest-bond approximation)[1,5]. It is possible that the amoeba redistributes its attachment molecules once a KV unit breaks. However, this will be a slower process compared to the failure process. Therefore, we can assume that the total force remains constant for a specific protrusion until it breaks completely. In addition, for the protrusion to remain intact until a length $L$, all KV units must survive. The survival probability of a single KV unit is

$$s_{\text{unit}}(L) = e^{-\int_0^L r(l)dl}. \tag{4}$$

Therefore, the survival probability of the whole protrusion is

$$S(L) = [s_{\text{unit}}(L)]^{n_0} = e^{-n_0 \int_0^L r(l)dl}. \tag{5}$$

We assume that the protrusion breaks when the survival probability becomes small, say $\tilde{s}$, a predefined constant. Thus, at a breaking length $L = L_b$, we have $S(L_b) = \tilde{s}$. Therefore,

$$e^{-[n_0 \int_0^{L_b} r(l)dl]} = \tilde{s}$$
$$\int_0^{L_b} r(l)dl = -\frac{ln(\tilde{s})}{n_0} \equiv \frac{s^*}{n_0}$$
$$r_0 \int_0^{L_b} e^{\alpha l} dl = \frac{s^*}{n_0}$$
$$L_b = \frac{1}{B} ln\left(1 + A/n_0\right) \tag{6}$$

where $A = \bar{\alpha} k s^* / \bar{r}_0 e^{\xi v}$ and $B = \bar{\alpha} k$. Thus, the theory predicts that thicker protrusions (with larger $n_0$) break at shorter lengths if the other parameters remain the same. As shown in Figs. 4(d) and (e) in the main text, this prediction agrees well with the experimental data. Note that both $A$ and $B$ depend on the mechanical parameters that will depend on the cellular state and other physical conditions such as the shape of the cell, leading to variability in the data.

Assuming that protrusions break at similar lengths, we can also compare the breaking time in the experiments and the theory. From Eq. (3), we can define a time scale $T_b$ from the scaled function when $(\ell(t) - \ell_0)/a$ has a certain value, say 80%. This gives us $T_b \sim 1/b$. We show in Table I below that the breaking times in the experiment and the theory agree well with each other.

TABLE I. Mechanical parameters of wild type SW480 and DLD1 cell lines.

| Avg. Parameter | WT SW480 | WT DLD1 |
|---|---|---|
| Breaking time, $T_b$ (s) | 21.59 | 36 |
| $b$ | 0.04 | 0.03 |
| $T_b = 1/b(s)$ [from theory] | 25.0 | 33.3 |

#### III. Properties of the Latrunculin-treated cells

We have also analyzed the data for Latrunculin-treated cells and found that they are well-described by the theory, Eq. (3). This means that the microscopic picture of the protrusion and the breaking mechanism remains the same. For example, Fig. 4(e) in the main text shows the $L_b$ vs thickness behavior, and it agrees well with the theory. Figure 1 shows that the experimental data collapse to a master curve, much like the WT data.

The breaking-times are similar for Latrunculin-treated cells ($\sim 65s$). Therefore, for better statistics, we have analyzed the data for both LatrSW480 and LatrDLD1 together. We show the comparison of breaking times for the average WT and Latr cell types in Table II.

$T_b$ from theory for Latrunculin-treated cells are slightly smaller than the experimental value,

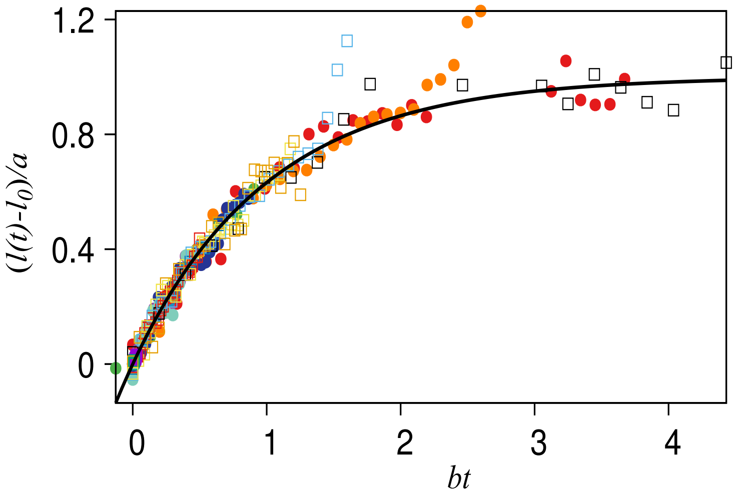


FIG. 1. Collapsed length vs time data of Latrunculin treated SW480 (circles) and DLD1 (box). Averaged $b = k/\xi = 0.02s^{-1}$.

TABLE II. Mechanical parameters of WT and Latrunculin-treated conditions. SW480 and DLD1 data are averaged.

| Avg. Parameter | WT | Latr |
|---|---|---|
| $T_b$ (s) [Experiment] | 28.79 | 65.73 |
| $b$ | 0.035 | 0.02 |
| $T_b = 1/b$ [Theory] | 28.6 | 50.0 |

the discrepancy might be due to lower statistics. However, the general trend is the same. The protrusion breaking time for Latrunculin-treated cells is nearly 2 times higher than that for WT cells (Table II). This explains why Latrunculin-treated cells show a smaller number of scission events in the experiment. Interestingly, although the Latrunculin-treated cells exhibit a fluid-like phenotype compared to WT cells, as evidenced from the AFM measurements, they show fewer trogocytic events and take more time for the scission. This trend is correctly captured by the theory.
